# Primate neocortex performs balanced sensory amplification

**DOI:** 10.1101/2022.06.23.497220

**Authors:** Jagruti J. Pattadkal, Boris V. Zemelman, Ila Fiete, Nicholas J. Priebe

## Abstract

Sensory cortex amplifies relevant features of external stimuli. This sensitivity and selectivity arise through the transformation of inputs by cortical circuitry. We characterize the circuit mechanisms and dynamics of cortical amplification by making large-scale simultaneous measurements of single cells in awake primates and by testing computational models. By comparing network activity in both driven and spontaneous states with models, we identify the circuit as operating in a regime of balanced amplification. Incoming inputs are strongly but transiently amplified by recurrent excitation. Inhibition acts to counterbalance this excitation by rapidly quenching responses, thereby permitting tracking of time-varying stimuli.

**One-Sentence Summary:** Sensory cortex uses balanced excitatory and inhibitory circuitry to boost weak signals while maintaining fast sensory dynamics in a changing environment.

## Main Text

Sensory cortical circuits are remarkable for their high sensitivity and precise selectivity for specific features in their inputs, even when these inputs are weak, noisy, and changing^1–5^. These properties are proposed to emerge from specific neuronal connectivity patterns within the cortex that act to amplify inputs and sculpt outputs^6–15^. We demonstrate how circuit organization can lead to distinct network dynamics. By combining a computational approach with empirical single cell activity measurements from large populations of neurons in awake primate area MT, a visual area where neurons are sensitive to visual motion ^16–19^, we uncover the operating regime of sensory cortex and supply fundamental constraints on its underlying circuitry.

To reveal the circuit origins of cortical amplification, we examined the dynamics of sensory neurons in visual area MT of awake primates. We recorded both spontaneous and evoked largescale population activity from inhibitory interneurons using two-photon imaging of calcium signals^20^ (Fig. 1A; 69 imaging sessions, 5046 neurons). Neurons in area MT were selective for motion direction^6,21,22^ and exhibited a functional map-like organization similar to primary visual cortex ^23^(Fig 1B,C). Direction tuning shifted systematically across the surface of area MT, such that nearby neurons had similar direction preferences (Fig. 1D). We characterized this organization by quantifying how the preferred directions of individual cells varied as a function of cell separation in cortical space (Fig 1E, length constant = 244 microns). Hypercolumn estimates across animals and chambers were similar (mean = 353 microns ± 129 s.d., Sup. Fig. 1A,B). This functional organization was stable when imaged over days or months (Sup. Fig 2).

**Fig. 1.**
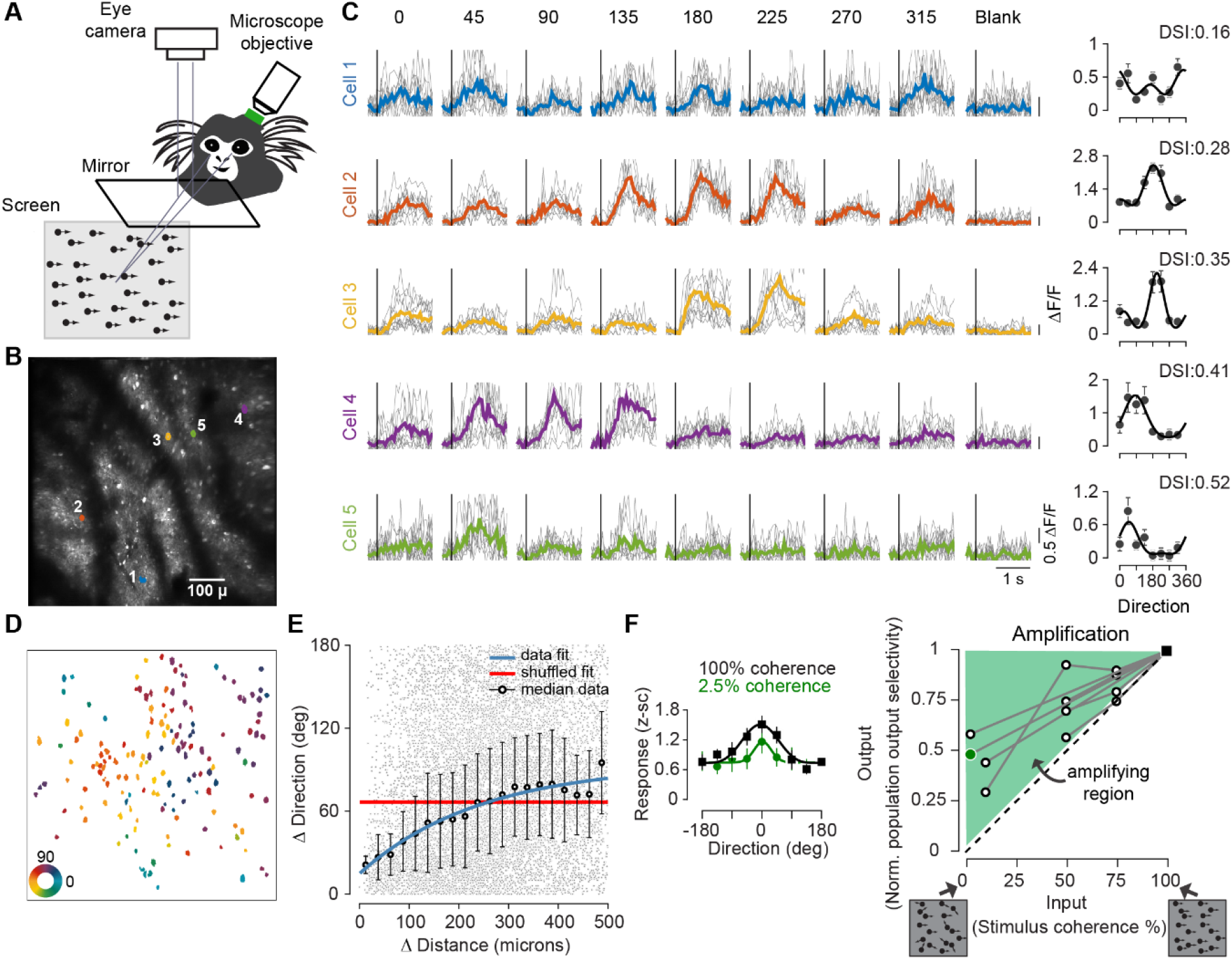
Population response characteristics in area MT. **(A)** Schematic of the experimental setup. The marmoset is head-fixed and sits in a chair in front of the display monitor, under the tilted objective of a two-photon microscope above area MT. An infrared camera allows eye tracking. **(B)** An example imaging plane in area MT. The locations of 5 example cells are highlighted. **(C)** Responses of the 5 example cells to fullfield motion in different directions. Gray lines show individual trial responses and colored lines are average response. Scale bars indicate 0.5 ΔF/F. Plots to the right are direction tuning curves. The plotted points are mean response per direction and error bars show the standard error. The responses were fitted with a Von Mises direction tuning curve. Each cell’s direction selectivity index (DSI) is indicated on top of the tuning curve. **(D)** Organization of preferred direction within plane. Outlines of cells with a DSI ≥ 0.15 from the same imaging field as in B are colored with their computed preferred direction. **(E)** Nearby cells share direction preferences. Difference in preferred direction between cells is plotted as a function of the physical distance between cells. Each dot is a cell pair. All cell pairs with DSI ≥ 0.15 are considered. The distances are then binned in 25 μ and the black circles represent the median direction difference for each bin. Error bars represent the angular standard deviation. Blue line is an exponential fit to the data and red line is an exponential fit to the shuffled data. **(F)** Amplification of direction signals. Responses of an MT population to high (100%) and low (2.5%) coherence motion, in which neurons are binned by direction preference (left panel) and fit by Von Mises direction tuning curves. The normalized selectivity of the population output, measured by the population DSI, is compared to the stimulus coherence (right panel). Lines connect responses from the same populations, different lines correspond to different populations. The green dot and black square represent the example population shown in left panel. Normalized selectivity is computed by dividing population DSI by the population DSI at 100% coherence. Even at low coherence, the population output is selective and the value lies above the diagonal, indicating amplification of input (shaded region).

Area MT neurons specifically amplify signals related to motion direction, even in the presence of high noise^4^. For example, MT population responses remain selective even when motion signals are degraded by lowering coherence to 2.5% (Fig. 1F).

To elucidate the circuit basis of this feature-specific amplification we integrated several recurrent amplification models^14,24,25^ into a unified computational framework (Fig. 2A; analytical derivations in Supplementary Information, Sup. Fig. 3). We explored non-normal networks, with segregated excitatory and inhibitory cell populations, in which two key model parameters alter network behavior: the strength of tuned recurrent excitation and the ratio of tuned recurrent excitation to tuned recurrent inhibition. The space of models that generate selective amplification can be divided into three main regions: one in which amplification emerges from balanced increases in excitation and inhibition and two in which excitation is dominant. Each of these networks has the same framework but varies in the strength of excitatory and inhibitory synaptic interactions between neurons (Fig. 2B), leading to different dynamics (Fig. 2C). Increases in tuned excitation lead to increases in amplification across all regions (Fig. 2D-F).

**Fig. 2.**
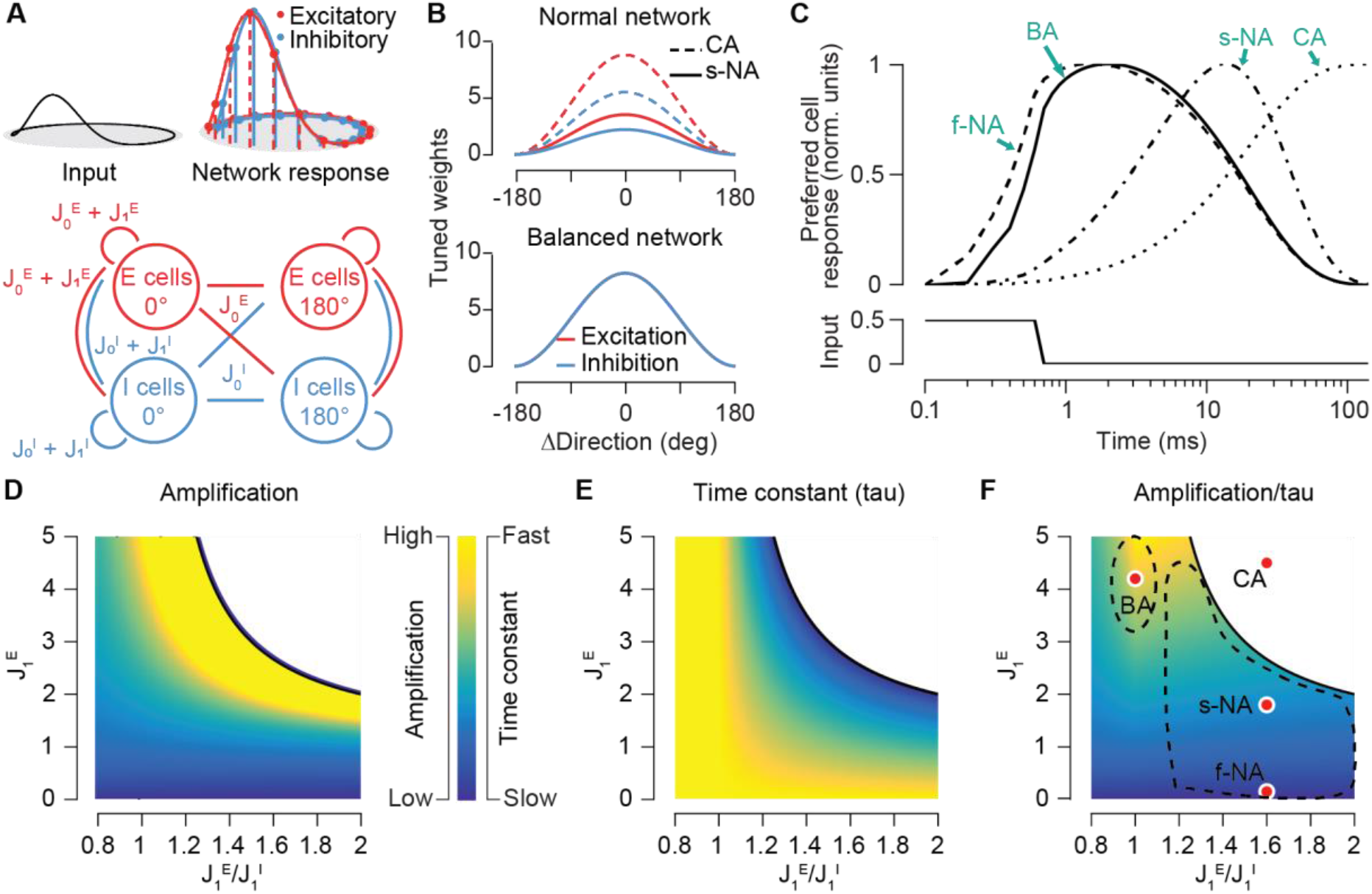
Circuits for selective amplification. **(A)** Top: Cartoon schematic of the cortical networks, including an input and separate excitatory and inhibitory populations. The input at direction θ is transformed by the model network to generate the population output shown at the top. Bottom: Cartoon schematic of connectivity parameters Weights J_0_^E,I^ refer to untuned weights from excitatory or inhibitory cells; weights J_1_^E,I^ are the tuned interactions. All cells are connected to each other with untuned weights, and both E and I cells interact with other cells of similar direction preference through the tuned weights, which vary by differences in direction preference (see B) **(B)** The tuned excitatory (red) and inhibitory (blue) tuned weights as a function of difference between cell preferences for direction. The top panel shows the excitatory and inhibitory tuned synaptic weights for CA (solid lines) and s-NA (dotted lines) networks. The bottom panel shows tuned synaptic weights in the BA regime, excitatory and inhibitory tuned weights are the same in this regime **(C)** Dynamics of responses in the model regimes are shown using responses to a brief step input (bottom). The f-NA (dashed line) and BA (continuous line) regimes responses are fast and decay rapidly, s-NA (dash dot line) responses are much slower and CA (dotted line) responses are persistent **(D-F)** Model behavior depends critically on the ratio of the tuned excitatory (J_1_^E^) and inhibitory (J_1_^I^) strength and the amplitude of the tuned excitatory component. The degree of amplification of motion inputs (D) 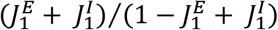, see analytical derivations), the inverse time constant (E) and the ratio of the amplification and time constant (F) are shown across model parameters. Red points indicate specific network parameters that are explored. The dotted region centered around BA indicates the balanced amplification zone. The bigger dotted region to the right of it indicates the normal amplification zone. States beyond the continuous curved line are in the continuous attractor regime. The boundary is defined as J_1_^E^ > (1+ J_1_^I^) (see analytical derivations).

Within the excitation dominant region, at low magnitudes of tuned excitation, the network’s baseline state is a single fixed point (normal amplification: NA), whereas at higher weight magnitudes, the network’s fixed points are a ring of tuned high activity states (continuous attractor: CA). Withdrawal of stimulus causes the network to decay to its baseline stable state in the NA regime^24^, with a slowing of dynamics as tuned excitation increases, until eventually in the CA regime the activity persists even in absence of the stimulus (Fig 2C,E). A sliver of the NA regime has fast dynamics but weak amplification (the f-NA regime); most of the NA zone exhibits larger amplification and slower dynamics (the s-NA regime). The balanced amplification (BA) regime consists of a circuit with strong tuned excitation and inhibition in a nearly matched configuration^26^. Like the NA regime, it exhibits a single fixed point at baseline activation. But unlike the NA and CA regimes, it generates strong amplification through a process of transient dynamics (Fig. 2C), in which small inputs along a feature dimension are selectively amplified, but then subsequently rapidly quenched by inhibition^25^. The BA regime only occurs in networks with segregated excitatory and inhibitory populations (Sup. Fig. 3). Increasing the strength of tuned excitation in the BA regime increases amplification, but unlike the NA regime, is not accompanied by a slowing of responses (Fig. 2D,E). Though these regimes are all selectively amplifying, they differ in the neural dynamics, which allows us to examine empirically the operating regime of cortical circuits.

We characterized MT dynamics at stimulus offset using a combination of intracellular and extracellular recordings. Networks in the CA regime should exhibit persistent activity even after stimulus removal, but MT cell activity rapidly declined to baseline after stimulus withdrawal, both at the level of membrane potential (Sup. Fig. 4A-D) and spike rate (Sup. Fig. 4E-H). These rapid offset dynamics are inconsistent with dynamics purely in the CA regime but are feasible with the BA or f-NA regime. They are also consistent with the slow amplifying s-NA regime: the decay to baseline stable state is rapid as it is not a part of the set of slow amplifying states. Finally, the dynamics are also possible in a version of NA regime (termed NA/CA) where a global increase in untuned input can shift the network from NA into a state where a ring attractor emerges (CA), while withdrawal of such an input rapidly shifts the network back ^27^. This NA/CA model with global input that is dynamically linked to the tuned stimulus could then exhibit rapid stimulus offset responses. Therefore, to distinguish between these alternatives, it is necessary to examine network dynamics when the stimulus is always present.

We next compared model and area MT network responses when stimulus is always present, but motion direction is rapidly switched (Fig. 3A). In the BA and f-NA regimes, network activity shifts abruptly with stimulus direction changes: the population activity bump induced by the initial motion direction decays rapidly in place while a new bump emerges to represent the second direction (Fig. 3B). In contrast, in the NA/CA networks, the network activity bump moves smoothly along the ring of directionally tuned responses, passing through and activating neurons selective for intermediate directions along the way (Fig. 3B, green traces). These model predictions do not change when the ratio of number of E cells to I cells is changed to 4:1, as observed in the neocortex (Sup. Fig. 5).

**Fig. 3.**
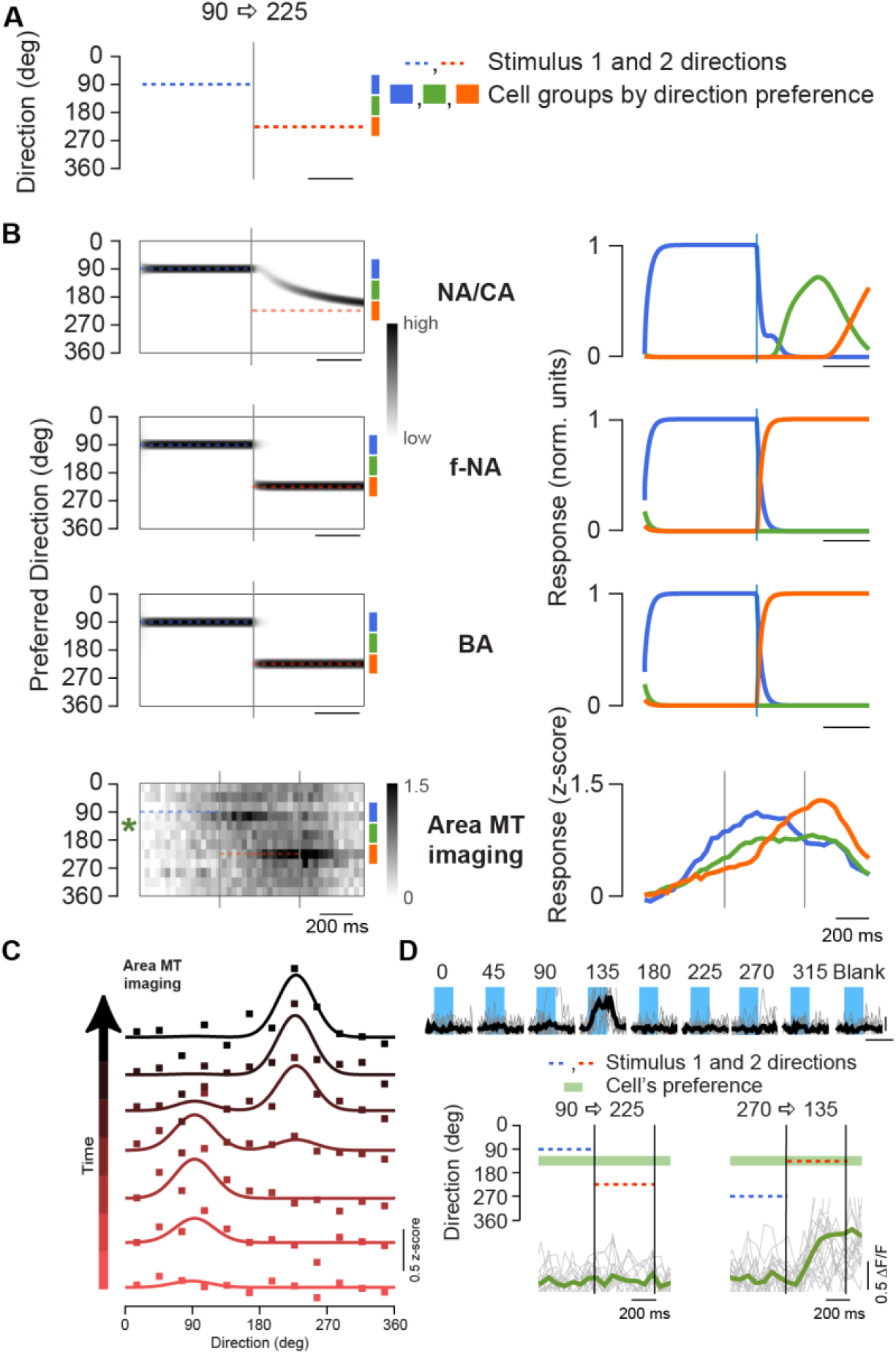
Shifts in motion direction elicit rapid changes in MT population responses. **(A)** A cartoon of the stimulus in which stimulus direction is rapidly changed from 90 degrees to 225 degrees. Blue and orange dashed lines indicate first and second stimulus intervals respectively. **(B)** Response of model networks to a sudden 90-225 direction change. Average response of a population of cells in a NA/CA network elicits a smooth rotation in the bump of activity from those cells preferring the first direction to the second direction (left). First stimulus starts at left edge of the heatmap for all panels. Next stimulus starts at the first gray vertical line. End of second stimulus is at the right end of model heatmaps. The end of second stimulus for calcium imaging heatmap is shown by the second vertical gray line. The response time courses of cells preferring the first direction (blue), the second direction (orange), or intermediate directions (green) are shown in the right panels. Both the f-NA and BA network populations exhibit rapid shifts (left panels), and neurons tuned to intermediate directions do not respond. Calcium recordings from area MT neurons also make rapid shifts in population activity, but there is a lag in the calcium imaged response relative to stimulus onset and offset due to the decay time of the calcium fluorescence signal. The intermediate cells do respond, but no more than expected from the response to the single stimulus conditions. **(C)** Population tuning curves at different epochs do not exhibit a smooth shift in preferred direction, but instead are well fit by a tuning curve with two modes in the first and second directions. **(D)** The response of a single cell selective for 135 degrees (top). This cell does not respond to a motion shift between 90 degrees and 225 degrees but does to a direction change between 270 and 135 degrees.

For a shift in stimulus direction from 90 to 225 degrees, the MT population response average shows two distinct activity bumps at the two presented directions, but no smooth transition between the bumps in the form of elevated activation of cells preferring intermediate directions (Fig. 3B, bottom). Splitting the population responses into three groups based on direction preference (those selective for the first, second or intermediate directions) reveals that the cells responded to the first and second stimuli with distinct latencies (Fig 3B, bottom right). Population tuning curves based on neuronal preference reveal that following the change in stimulus direction, a new bump appeared at the second direction while the activity at the first direction decayed (Fig 3C). Individual MT neurons tuned to intermediate directions responded weakly, not exceeding the amount expected based on their direct evoked response to the two motion directions, inconsistent with a bump of activation that passes through the intermediate directions (Fig. 3D, Sup. Fig. 6). The slow calcium signal may not fully register a rapidly moving bump that passes through the intermediate directions, but extracellular single cell electrophysiology measurements in MT revealed a similar absence of responses in neurons tuned to intermediate directions (Sup. Fig 6A-G). As predicted by our model, excitatory cells (from electrophysiology) and inhibitory cells (from imaging) exhibit similar population behavior.

To determine quantitatively whether MT population responses discretely switch their signals for direction (as in the f-NA/BA networks) or sweep smoothly through intervening directions (NA/ CA networks), we compared physiological network responses to switch and sweep predictions (Fig. 3C, Sup. Fig. 7). Both our calcium and electrophysiology records are highly correlated with a switch prediction and not the sweep prediction (Ca: R_switch_ mean = 0.52 +/- 0.2, R_sweep_ mean =0.13 +/- 0.09. Ephys: R_switch_ = 0.91, R_sweep_ = 0.13, Sup. Fig. 7B). One caveat to our interpretation is that the stimulus might be so strong that it overrides the slow internal dynamics of the NA/CA response. We therefore repeated the experiment at the lower contrast of 8% (Sup. Fig. 6H-J). Even at this contrast, we observed no evidence of a bump of activation moving through intermediate directions, indicating that area MT is not operating in the NA/CA amplification regimes.

Another distinguishing feature of amplifying models is how they respond to smooth decrements in signal strength (Sup. Fig 8A-B). In the NA/CA networks, an initial strong stimulus pushes the networks onto a stable ring and lowering the stimulus strength smoothly has little impact on network activity (Sup. Fig. 8C). In contrast, the f-NA and BA networks are sensitive to decreases in the input strength, showing smooth and large reductions in tuned response amplitude (Sup. Fig. 8C, bottom panels). Like the f-NA and BA networks, the tuned responses of area MT populations declined as coherence decreased (Sup. Fig. 8D-G). MT cell responses to changes in direction and coherence show that the network is highly sensitive to the amplitude and direction of the stimulus, inconsistent with operation in the NA/CA regimes but consistent with the BA and the f-NA regimes.

These candidate amplification models also exhibited distinct spontaneous activity signatures. When only noise input is provided the f-NA network remained nearly quiescent, with little tuned response. Networks in the balanced regime, however, exhibited large spontaneous activity fluctuations that are tuned, with varying amplitude and directionality. In contrast, NA/CA networks exhibited tuned responses persistently, even in the absence of structured input. Though spontaneous activity in MT is weaker on average than visually-evoked responses, it is characterized by sparse punctate bursts, with individual MT neuron amplitudes approaching those of evoked responses (Fig. 4A,E). The sparse activity events generated skewed activity distributions for both individual cell and network activity (Sup. Fig. 9A-D, skewness: 1.4, median skew =1.2 ± 0.5 s.d., across all sessions), a common phenomenon in neural circuits^28,29^. Cells of similar direction preference were co-activated during these spontaneous bouts of activity (Sup. Fig. 10), demonstrating that internally-generated feature-specific amplification occurs even in the absence of a motion stimulus^30–35^.

**Fig. 4.**
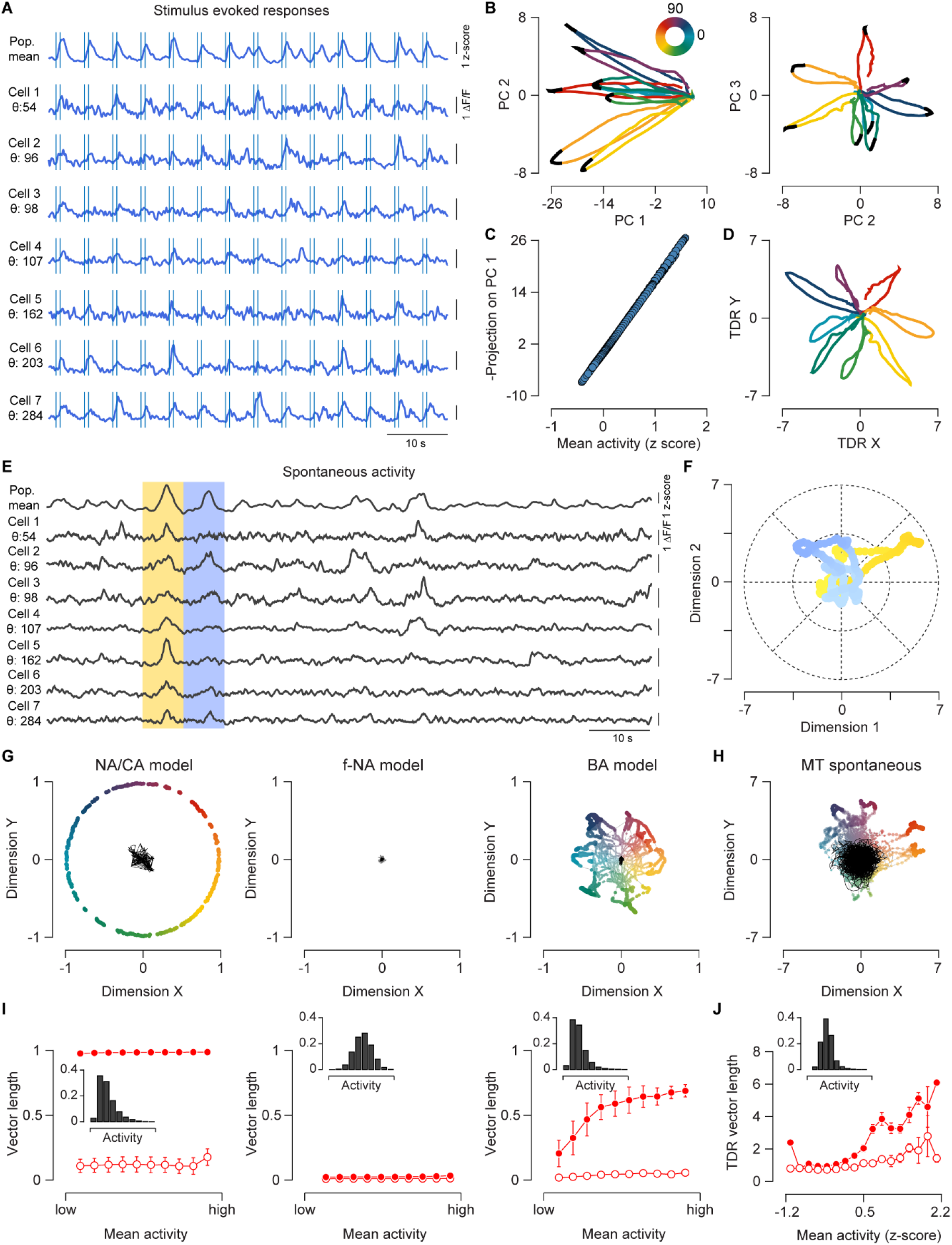
Area MT spontaneous and evoked activity analysis to differentiate between different circuit models. **(A)** Example of simultaneous stimulus-evoked activity traces across area MT neurons. The population mean z-scored cell activity (top row) is displayed above a subset of the neuronal population (bottom rows). The vertical lines indicate start and stop of the stimulus, motion duration was 700 ms. Vertical scale bars to the right indicate a 1 ΔF/F. The preferred direction of each cell is indicated by θ. **(B)** Stimulus-evoked population response trajectories in principal components space. Mean population response trajectory for each of the 8 directions presented are projected along the principal component dimensions. The color coding indicates the stimulus direction. The black region in each trajectory shows the peak response interval. The individual neuron contributions to PC2 and PC3 were dependent on motion selectivity with cells having a higher degree of motion selectivity contributing more to these PCs (linear correlation coefficient between DSI and absolute PC2 weights is 0.56, with PC3 weights was 0.25, Sup. Fig. 11B, C, top). The PC2 and PC3 weights also had a sinusoidal relation with the preferred directions of cells (Sup. Fig. 11B, C, bottom). **(C)** PC 1 is related to mean population response. The individual neuron contribution to PC1 did not depend on its degree of motion selectivity or direction preference (linear correlation coefficient between PC1 weights and DSI was 0.17 and between PC1 weights and preferred direction was −0.05, Sup. Fig. 11A). **(D)** Population response projections in TDR space. Stimulus-evoked mean responses are projected in TDR space and show a better separation along different directions. **(E)** Example of simultaneous spontaneous activity traces across the same area MT population as in A-D. The population mean of z-scored responses across all cells is shown at the top. Vertical scale bars indicate a response of 1 ΔF/F. The yellow and blue regions mark two active epochs. **(F)** Population trajectory during the two spontaneous epochs. Activity during the two epochs marked in blue and yellow is projected in TDR space shown in panel D. Hue corresponds to the activity, darker hue indicating higher activity. **(G)** Spontaneous activity predictions for the three circuit models. The simulated spontaneous activity is projected in direction space for the NA/CA network (left panel), which has responses lying on the attractor ring, the f-NA network, which lacks directional responses (middle panel), and the BA network, which shows varying directionality of the responses (right panel). The colors indicate the output direction of the population response. The size and opacity of the color-coded dots indicate activity amplitude with bigger dots and higher opacity for higher activity. Shown in black is the projection of the same data after shuffling direction preferences of model neurons. **(H)** The projection of MT spontaneous data into the TDR direction space indicates that there are significant deviations of the network response which signal direction spontaneously. Color-coded trajectories are the observed data and black is the shuffled responses. **(I)** Relationship between output directionality and activity in model networks. Spontaneous model activity and shuffled model activity are shown in filled and open circles, respectively. Error bars indicate the standard deviation. The inset histograms are normalized distributions of overall activity within the networks. **(J)** As in I, relationship of spontaneous activity projection to activity magnitude for the example area MT population. Error bars are 95% bootstrapped confidence intervals over the median. The normalized distribution of mean activity across all cells in spontaneous period is shown in the inset.

To compare quantitatively the patterns of spontaneous activity in the MT population with spontaneous model predictions, we projected the high-dimensional spontaneous population activity into a low-dimensional direction space. We calculated this space using evoked responses to motion stimuli. Motion stimuli evoked large increases in the overall population response, but the motion direction modulated which population subsets responded (Fig 4A). To visualize the tuning of the population response, we first performed unsupervised dimensionality reduction using principal components analysis (PCA) to generate a low-dimensional projection of the network activity. PCA provided a compact representation of the 260 recorded neurons described in Figure 4 as only 5 dimensions accounted for over 90% of the variance in the population activity. The dimensions that emerged from PCA were easily interpretable: the first dimension (PC1) corresponded to mean population activity (PC1, mean population activity regression R^2^= 0.98, fig 4B,C), the next two dimensions (PC2, PC3) encoded the motion direction, with adjacent motion directions encoded by nearby polar angles in the PC2, PC3 plane (Fig. 4B, Sup. Fig. 11). We consistently observed similar evoked response trajectories across multiple sessions in different animals (Sup. Fig. 12). PCA can provide a quantitatively distorted view of the population code for direction if the recorded sample contains unequal numbers of cells encoding different motion directions (Fig 4B). To correct for these distortions we applied targeted dimensionality reduction (TDR ^36^), which explicitly identifies the dimensions that capture most of the direction-related variance in the population. TDR provided a similar representation to the PCA but separated the directional responses better (Fig. 4D). Projecting spontaneous MT population activity into the evoked PC direction space revealed that the large spontaneous events in MT neurons resemble directionally tuned evoked responses (Fig. 4F), consistent with the concerted activation of neurons with common motion preferences (Sup. Fig. 10^30–35^). In rodent neocortex, which lacks a functional organization, the relationship between spontaneous and evoked activity may be distinct and depend on modality ^32,37^. The dimensionality of the spontaneous population response is higher than evoked, as 142 dimensions are required to explain 90% of the variance of the spontaneous activity. The first three PCs or the 1^st^ PC and the 2 TDR dimensions (Fig. 4F), however, accounted for a substantial fraction of spontaneous activity variance (23% and 22% respectively), similar to the fraction of the evoked response variance explained by these components (31% for both).

Comparing the projections of MT spontaneous activity into the low-dimensional direction space to the model predictions revealed a variable and fluctuating degree of directionality (spontaneous session data: Fig 4H, evoked session data: Fig 4D), consistent with the BA regime (Fig. 4G). In contrast to other network regimes, the degree of directionality in the BA network covaried strongly with the amplitude of network activity (Fig. 4I) and the network exhibited skewed population activity (skewness: 1.71). Notably, MT activity in the spontaneous state also exhibited a similar monotonic covariation between activity level and directionality. This covariation between the directionality and magnitude of activity was not explained by a global rescaling of low amplitude responses (Fig 4J) and was observed across multiple sessions in two animals (Sup. Fig 13), for both spontaneous and evoked responses.

We have used records from large scale populations of neurons and computational models to constrain the circuit mechanisms underlying selective amplification in sensory cortex. Our findings demonstrate that evoked and spontaneous cortical dynamics are consistent with balanced amplification while excluding the other potential networks. Cortical dynamics in MT are highly sensitive to time-varying sensory signals, displaying selective amplification and fast responses to changes in the inputs, properties that are well-modeled by a circuit operating in a balanced amplification regime. Cortical dynamics exhibit large tuned spontaneous activity fluctuations that resemble evoked responses. Finally, the degree of selectivity in spontaneous activity grows proportionally with activity. This combination of features is present only in a balanced amplification cortical circuit ^25,38–42^. Such a network achieves a balance between rapidly representing feedforward signals while boosting weak signals through recurrent connections (Fig. 2F). Note that our exploration of the cortical operating regime is in awake primate and this regime may change with differences in animal state such as attention and motivation ^43–46^.

It is interesting that a balanced directionally tuned circuit has qualitatively the same architecture as the mammalian head direction (HD) ring attractor circuit. Yet, strikingly, they operate in completely different regimes that are rooted in the distinct nature of the computations that must be performed: rapid balanced amplification of inputs in sensory cortex versus the storage of a persistent memory of head direction and integration of head velocity signals through continuous attractor dynamics in the HD system ^47^. Though only relatively subtle adjustments in the relative strengths of excitatory and inhibitory connectivity are necessary to shift a network from a balanced amplification regime with a single attractor, as observed in area MT, to a ring attractor (Fig. 2), as observed in the HD circuit, the resulting changes in dynamics and function are large.

The balanced amplification regime only emerges from models in which there are distinct excitatory and inhibitory neurons (Sup. fig 3,^25^). That sensory cortex operates in the balanced amplification regime provides a potential explanation for the specialization of neurons into distinct excitatory and inhibitory populations: the essential asymmetry that arises from coupling these populations is fundamental to the generation of large but responsive selective amplification of time-varying stimuli.

## Supplementary Materials

Figs. S1 to S13

## Methods

Analytical model derivations and description

## Acknowledgements

We would like to thank David Hansel, Robbe Goris and Andrew Tan for their comments on the manuscript. We also thank Allison Laudano and Carrie Barr for assistance with animal care.

## Author Contributions

Conceptualization: JJP, IF, NJP

Methodology: JJP, BVZ, NJP

Investigation: JJP, NJP

Visualization: JJP, IF, NJP

Funding acquisition: BVZ, IF, NJP

Project administration: BVZ, IF, NJP

Supervision: BVZ, IF, NJP

Writing – original draft: JJP, IF, NJP

Writing – review & editing: JJP, BVZ, IF, NJP

## Competing interests

None

**Supplementary figure 1:**
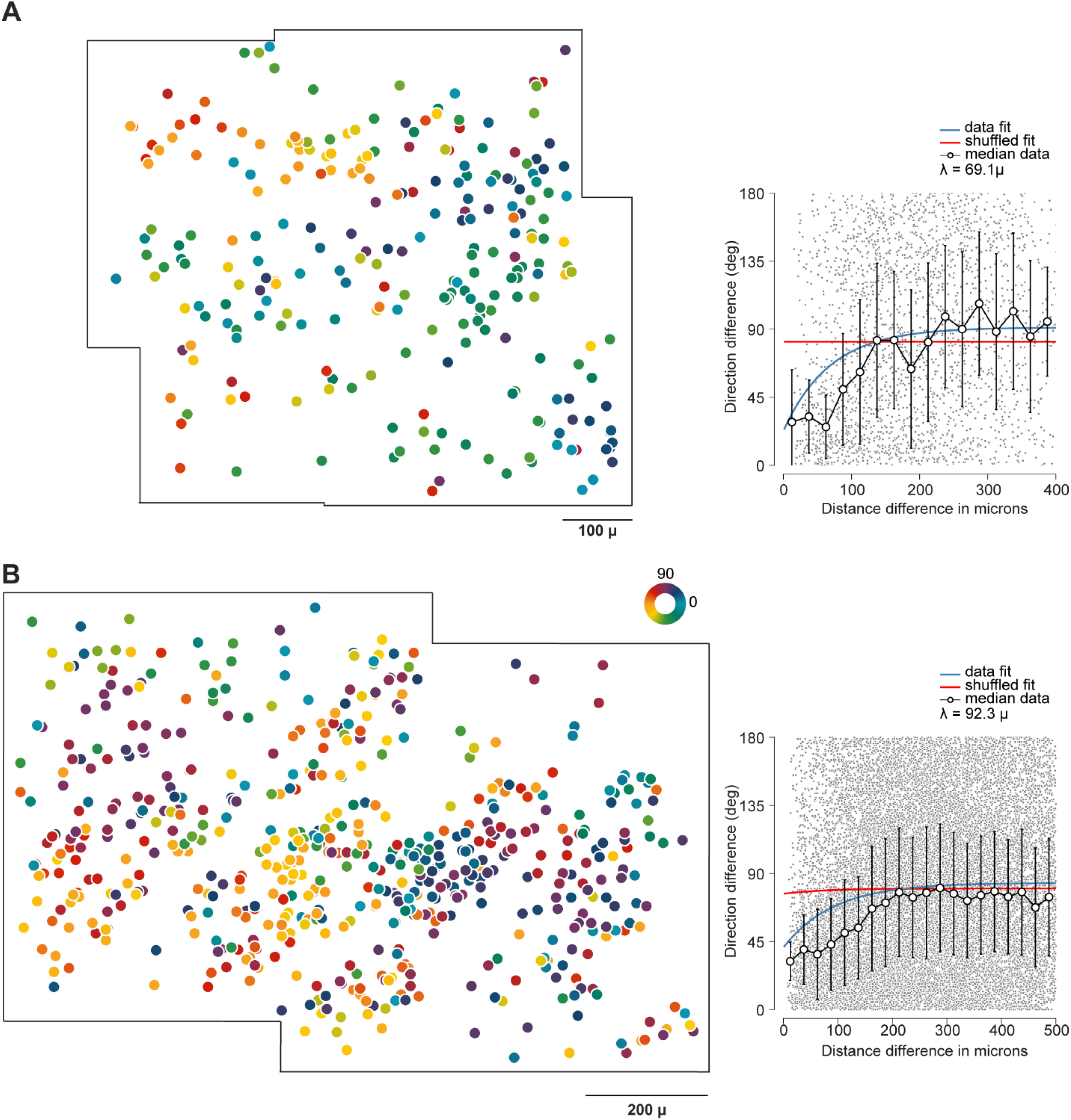
Example area MT direction maps collected on single days. A) A direction map generated by pooling across planes (left panel). Each cell is denoted by one dot and colored by its direction preference (see color wheel). The pairwise dependence of difference in cortical distance and direction preference for this map is shown in the right panel. B) Direction map measured from another chamber.

**Supplementary figure 2:**
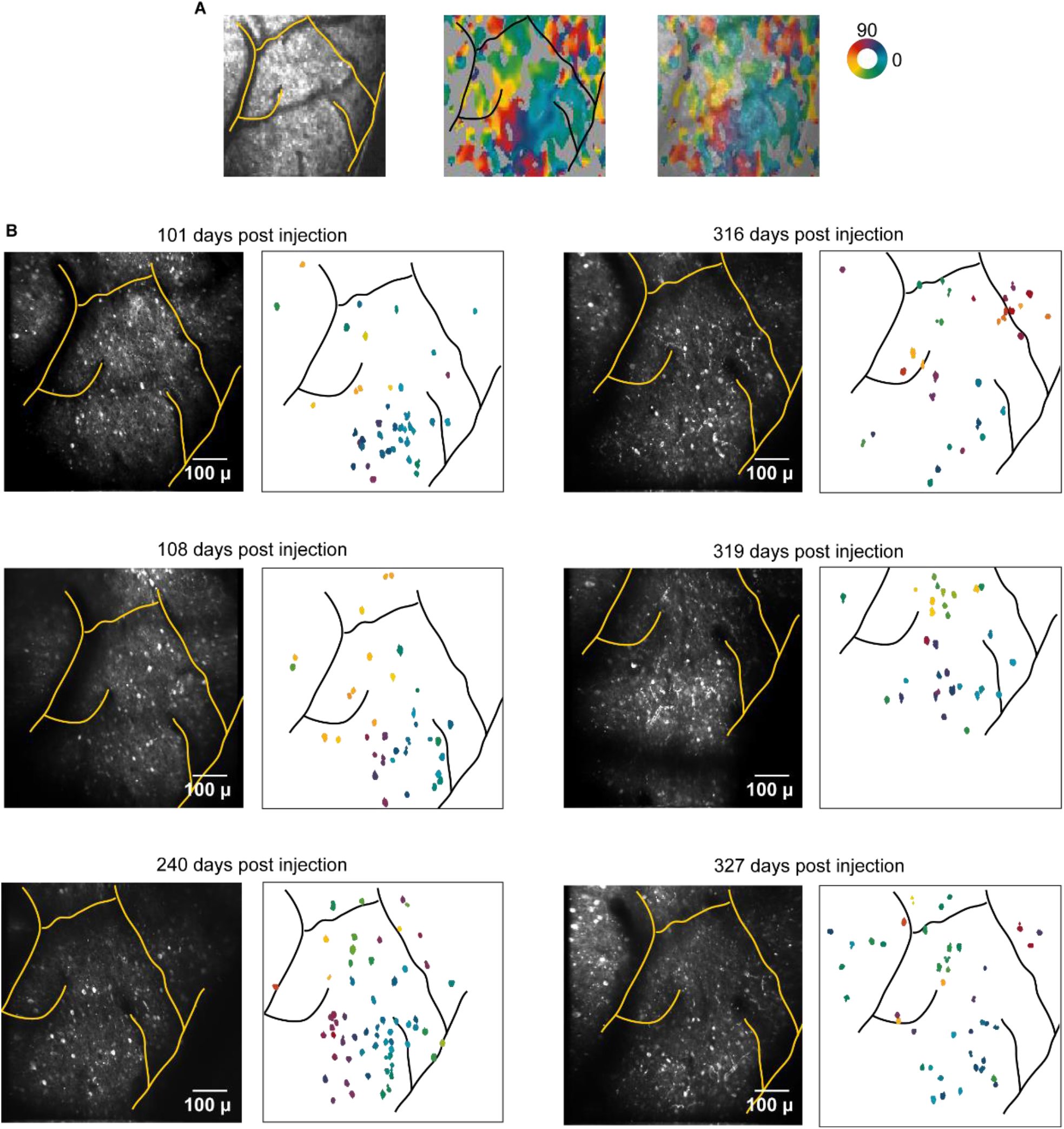
Stability of the functional organization. A) Wide-field microscopy provides images of the vasculature at an example site (left panel) and the direction map of the same site (middle panel). These images are overlaid in the right panel. B) The same area is imaged across multiple days at cellular resolution. Each panel indicates the days past the virus injection. Within each panel, the imaging plane (left) and the organization of direction selective cells obtained from that side (right) are shown. Each colored area is a cell color coded by its direction preference. Outlines of the vasculature are drawn over each area as the precise imaging location varies.

**Supplementary figure 3:**
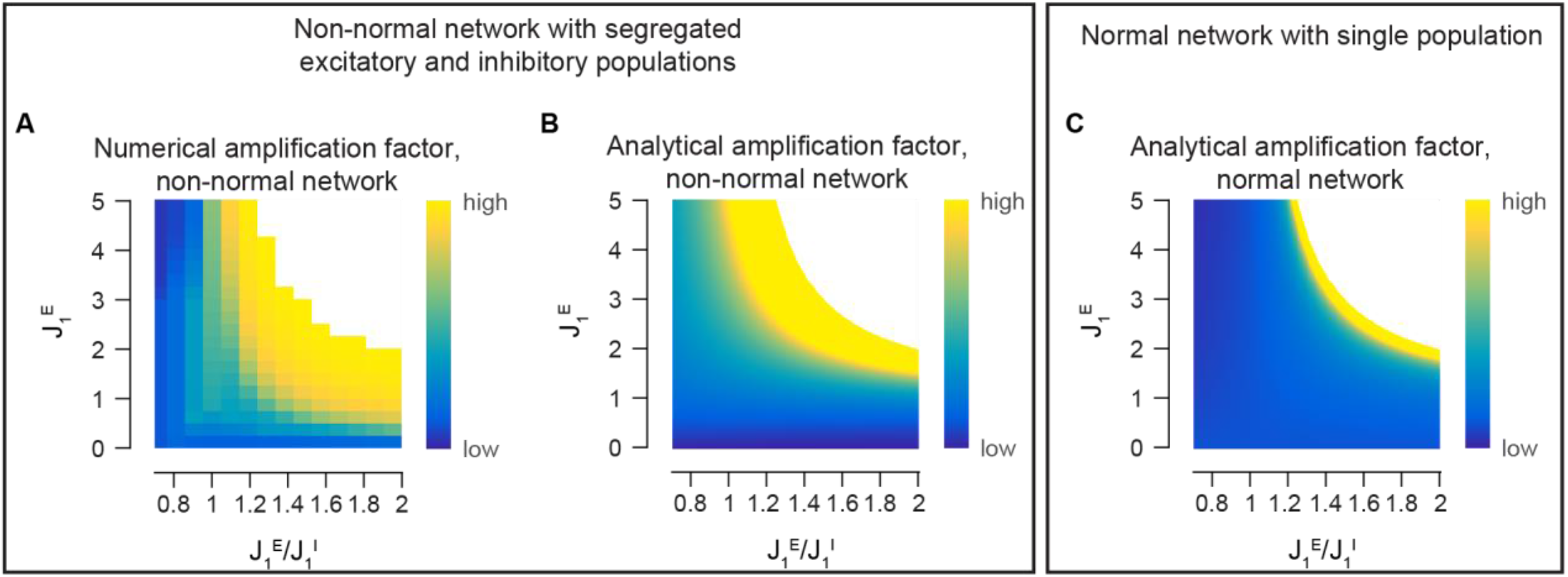
Amplification factor comparisons across non-normal and normal networks. A) Amplification factor for numerical simulations with non-normal network. B) Analytical amplification factor for non-normal network. C) Analytical amplification factor for normal network, without segregated E and I populations.

**Supplementary figure 4:**
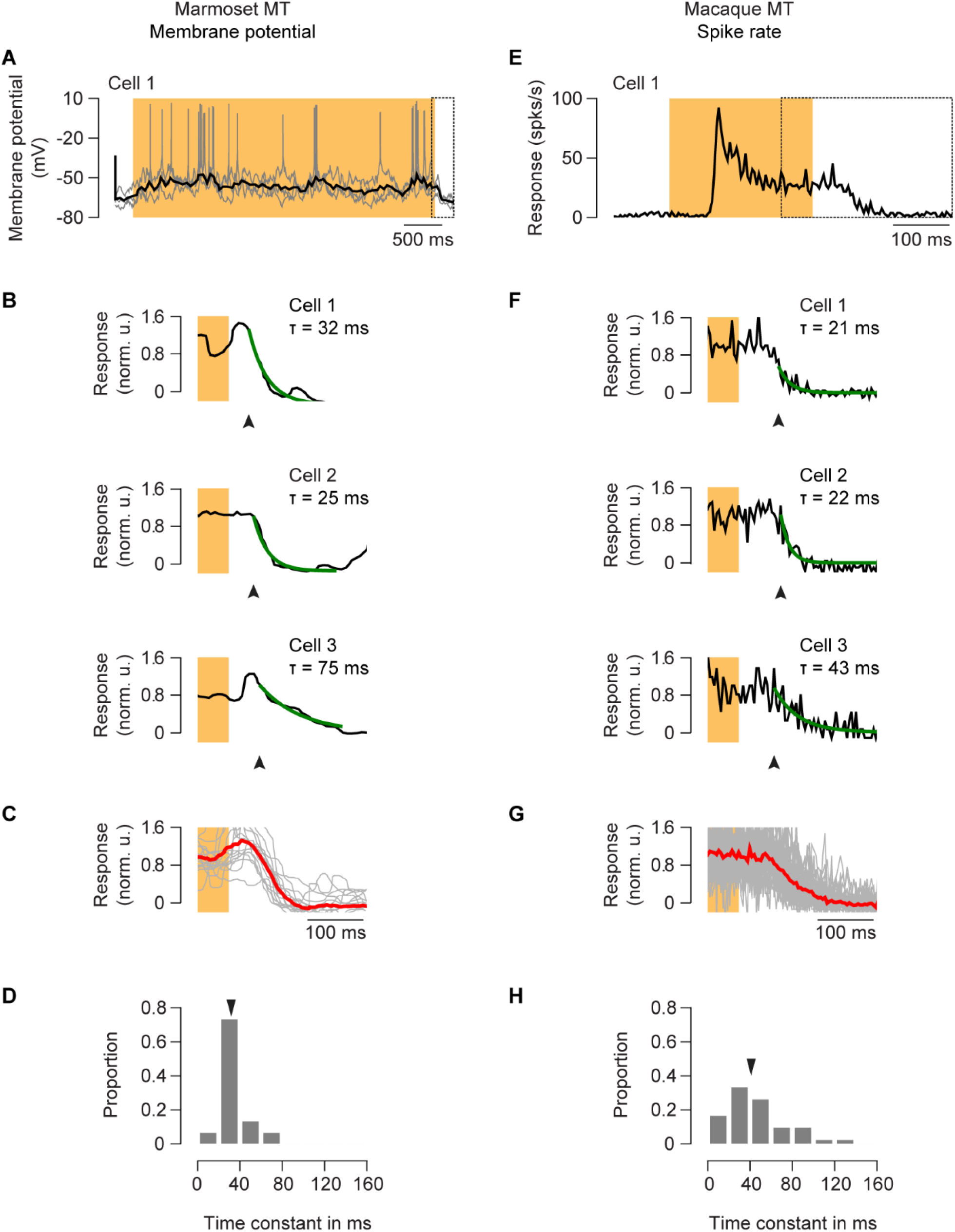
Offset responses in area MT measured with intracellular and extracellular recording. A) Example cell membrane potential trace in response to stimulus motion recorded intracellularly from marmoset MT. Gray lines represent individual trials. Thick black line is the median filtered mean response. Box at the end of the trace shows the interval displayed in B. B) Whole cell records at the termination of preferred direction motion in marmoset area MT (cell 1 same as A). Lines indicate an exponential fit of the Vm decline to baseline activation. Responses are normalized by the mean depolarization induced by the visual stimulus. C) Average response at responses across neurons (red) and the individual neurons in the sample population (gray lines). D) The distribution of tau fit from the exponential decay to baseline. E-G) As in A-D, for extracellular recordings in macaque area MT.

**Supplementary figure 5:**
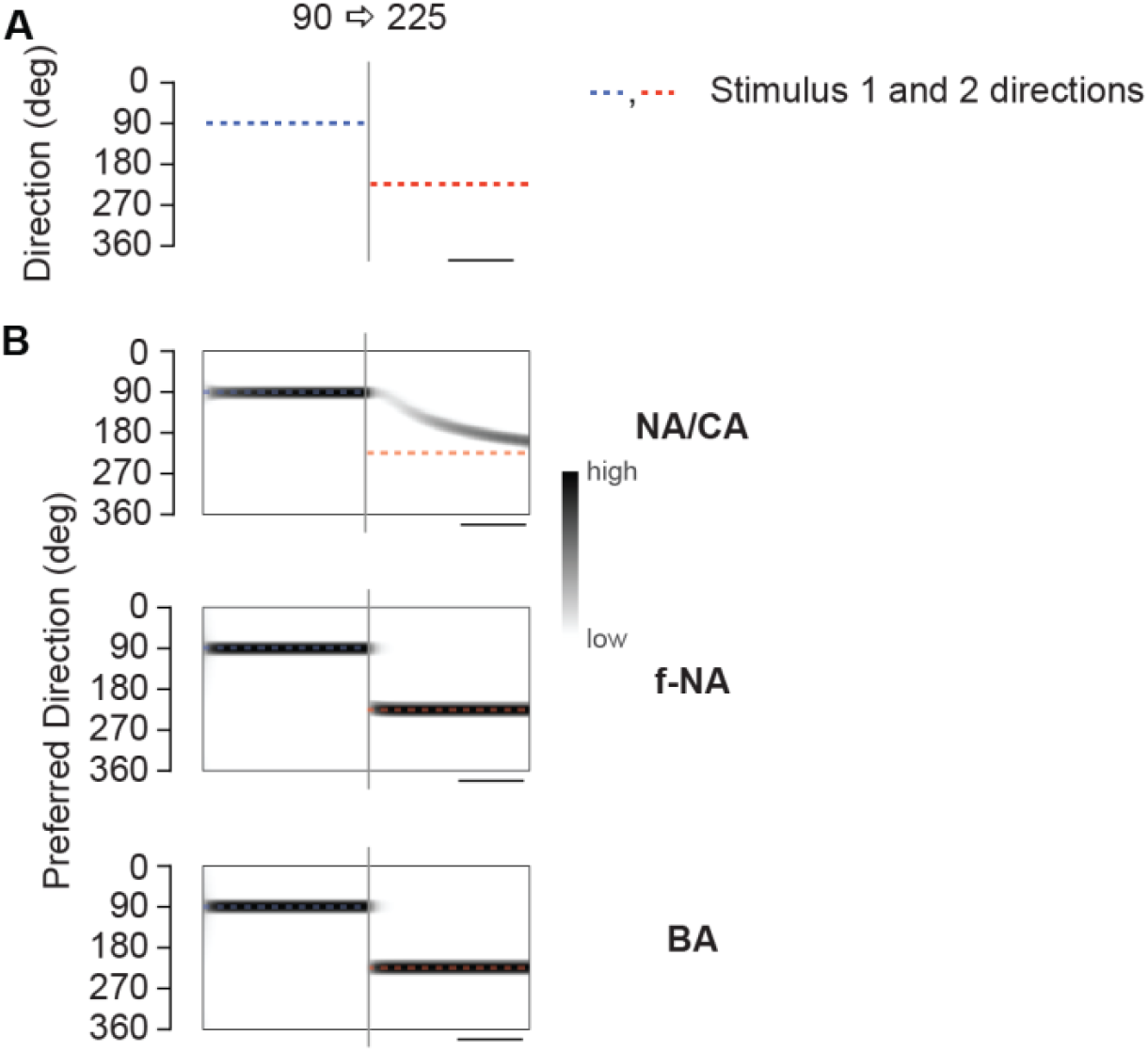
Model predictions for shifts in motion direction for E:I number of cells ratio of 4:1. (A) A cartoon of the stimulus in which stimulus direction is rapidly changed from 90 degrees to 225 degrees. Blue and orange dashed lines indicate first and second stimulus intervals respectively. (B) Response of model networks to a sudden 90-225 direction change. Average response of a population of cells in a NA/CA network elicits a smooth rotation in the bump of activity from those cells preferring the first direction to the second direction (left). First stimulus starts at left edge of the heatmap for all panels. Next stimulus starts at the first gray vertical line. End of second stimulus is at the right end of model heatmaps. Both the f-NA and BA network populations exhibit rapid shifts (left panels), and neurons tuned to intermediate directions do not respond. The model population has 4 times more E cells than I cells, the effective excitation and inhibition is held constant by modifying the weights.

**Supplementary figure 6:**
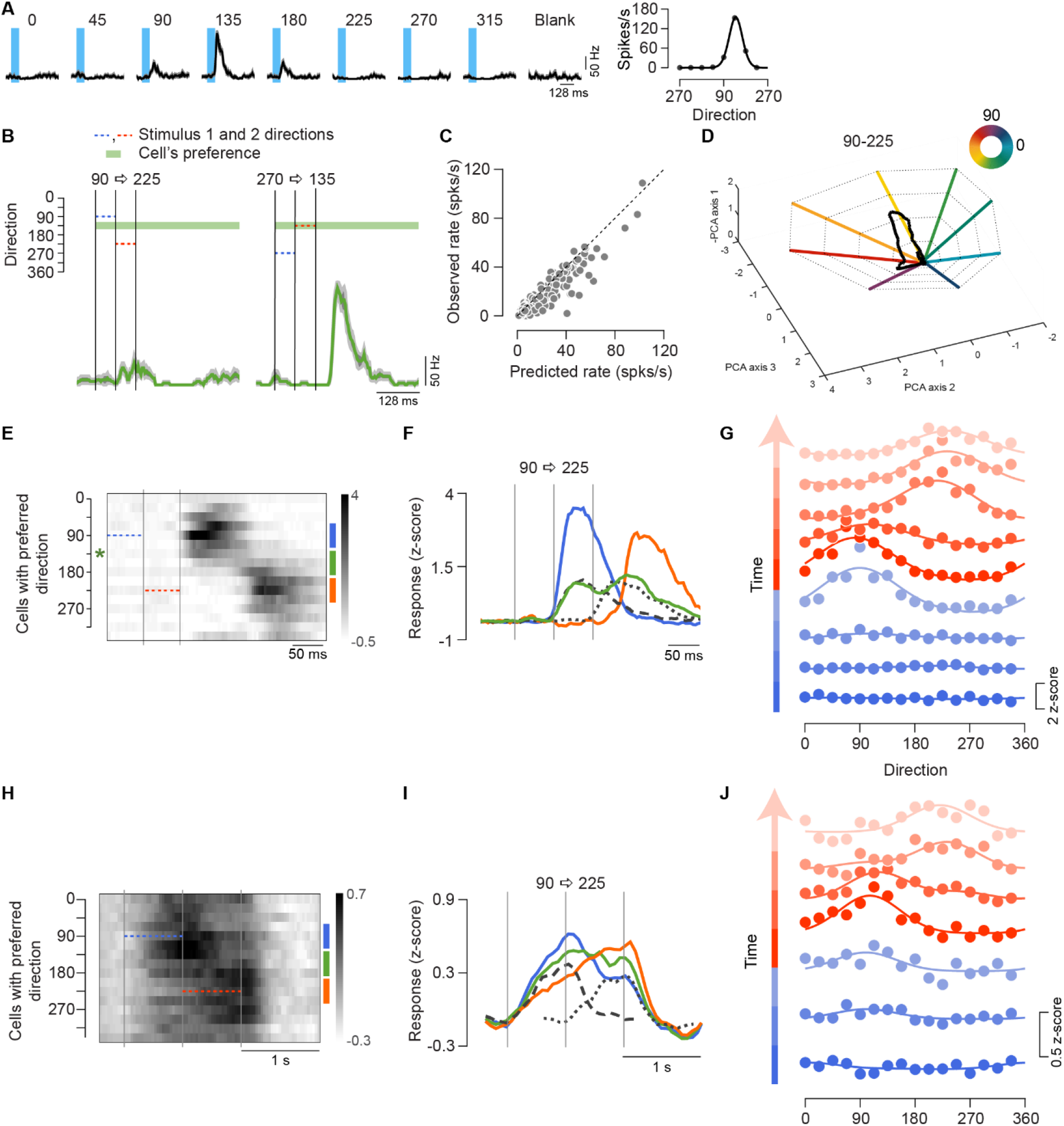
Rapid shift in MT responses for sudden change of stimulus direction using electrophysiology. A) Responses of an example cell for different motion directions. Its direction tuning curve is shown to the right. Cell prefers 135 degrees. B) Response of the example cell for 90-225 stimulus change. Blue and orange indicate first and second stimulus interval respectively (left). Response of the same cell for 270-135 stimulus change (right). C) Comparison of observed responses of all cells to 90-225 stimulus change to predicted responses based on tuning properties. See methods for predicted response estimation. Observed responses are significantly lower than predicted (t-test p-value: 5.5×10^−15^). D) Population trajectory in TDR space for 90-225 stimulus change. Thick lines indicate response directions for stimulus motion along color coded directions. Contours connect same response levels across directions. Population trajectory does not move along the ring, but directly changes from 90 (purple) to 225 (yellow). E) Average responses across conditions. Two discrete activity bumps are seen. F) Time course of three cell classes for 135 degrees stimulus change. Blue is average of cells preferring first presented direction, orange is average of cells preferring second motion direction, green is average of cells preferring intermediate direction. Dashed line is the response of the green cell class to stimulus 1 presentation alone, dotted line is response of the green cell class to stimulus 2 presentation alone. G) Population tuning curves at different time points. Slices of population output are shown at 8 ms intervals from start of stimulus 1. Activity bump transitions discretely from stimulus 1 to 2. H-J) As in E-G, but for calcium imaging using low contrast (8%) stimulus.

**Supplementary figure 7:**
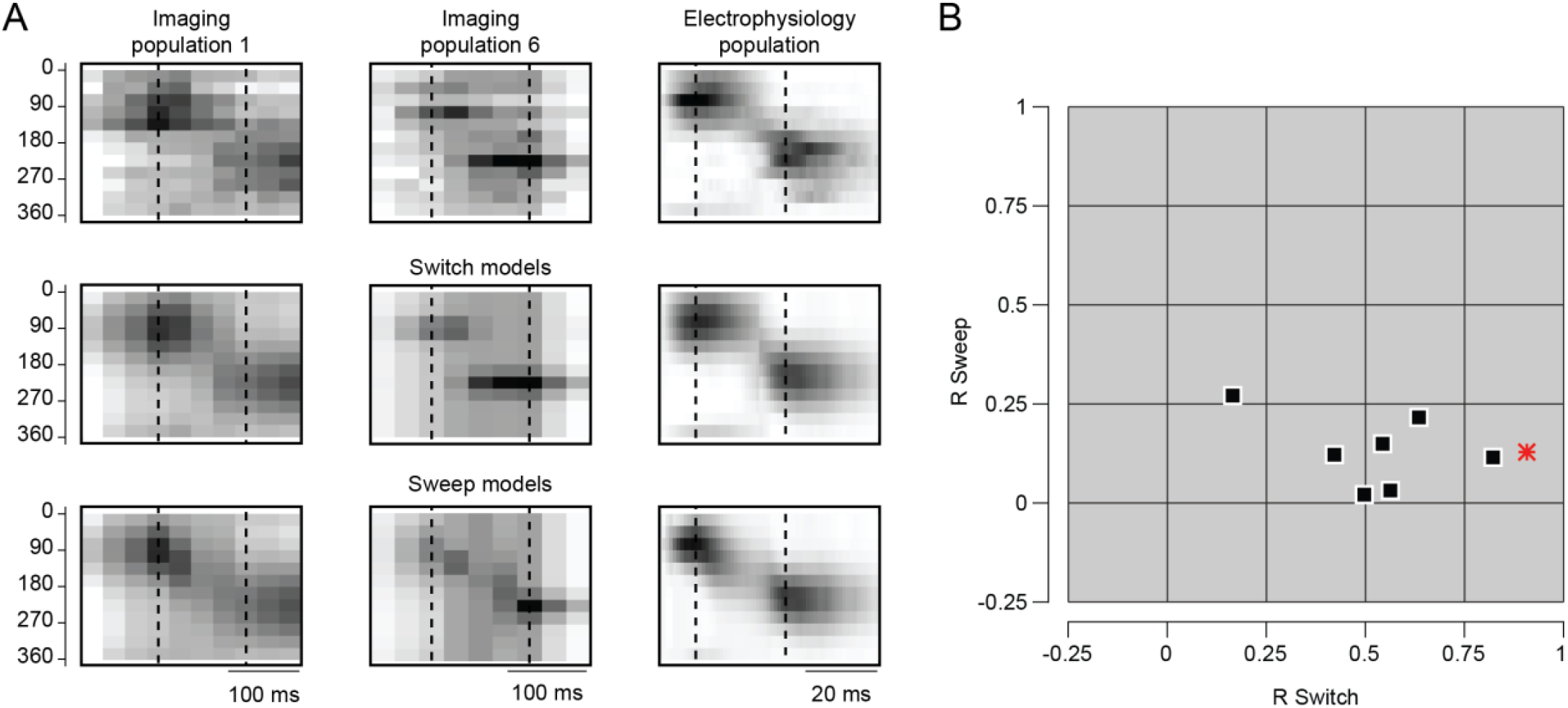
Direction changes induce switches in tuning, not sweeps. A) Example population responses are shown for 2 calcium imaging populations (left, middle panels) and one electrophysiology population. Data are shown in the top row. Switch model fits to the data are shown in the second row, in which the data is fit by population tuning curves for which the response is the sum of two Gaussian tuning curves separated by 135 degrees. In the bottom row the data are fit by a sweep model in which the population output preferred direction smoothly shifts from the first direction to the second direction. B) The partial correlation between the data and the switch model (abscissa) and the data and the rotation model (ordinate) are plotted. Each symbol indicates a distinct imaging session. The electrophysiology population is indicated by the red star.

**Supplementary figure 8:**
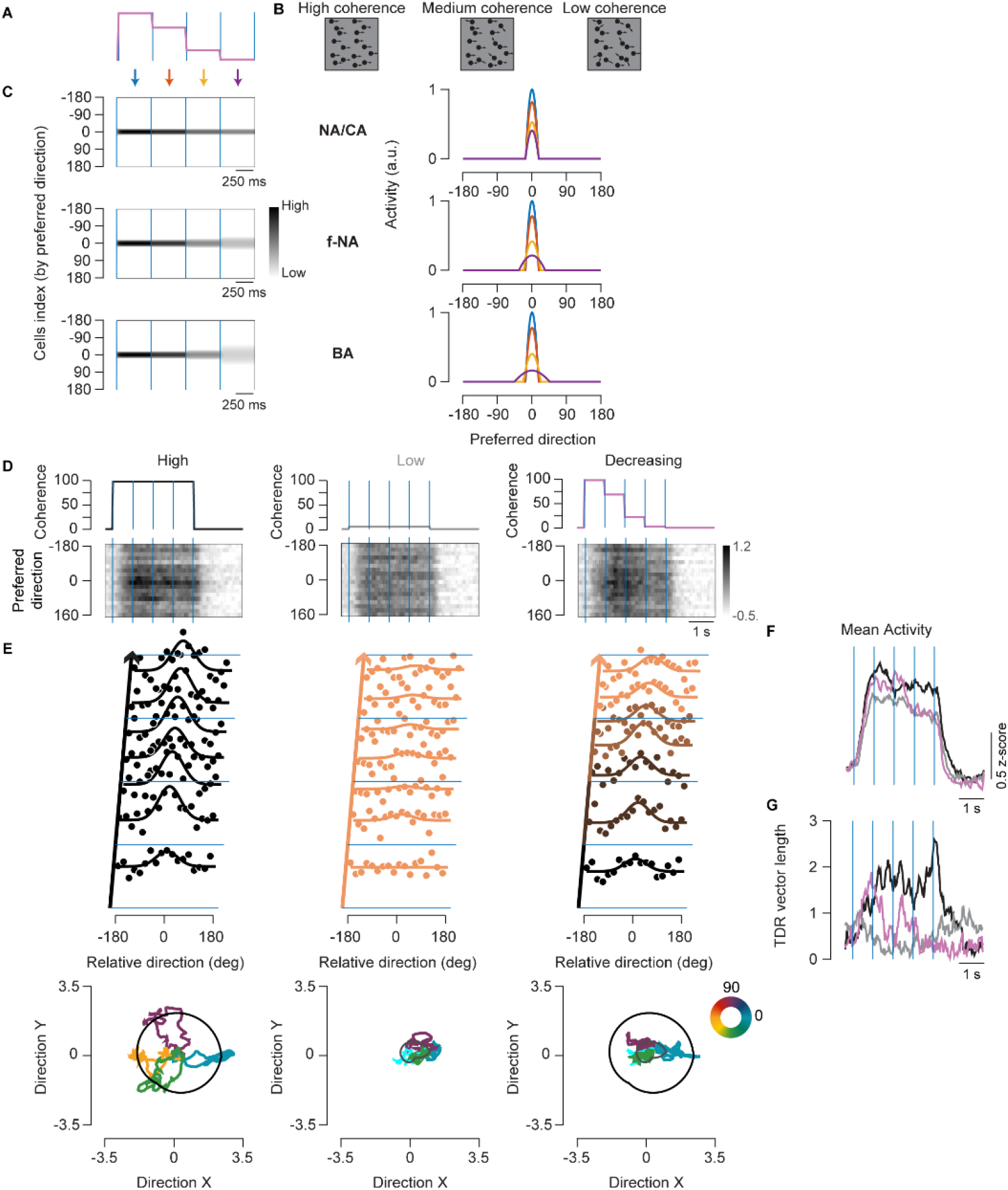
Network responses to changes in coherence. A) Population response prediction under s-NA/CAM, f-NA and BA for change coherence stimulus. Top schematic shows the change coherence stimulus. Each trial consists of 4 parts. The coherence value is constant in each part and changes in each part from high to low from beginning to end of trial. B) Shows the schematic for different coherence values. C) Population responses predicted in s-NA/CA, f-NA and BA networks are shown to the left. The s-NA/CA network maintains a near consistent tuning of population output even upon reducing coherence, whereas the f-NA and BA network output becomes less selective with decreasing coherence. To the right, the population output of the 4 coherence segments in the model networks for the model networks is shown. Color coding of segments is indicated in A. D) Population average across conditions. Neural responses are aligned such that stimulus is always at 0 degrees (bottom). The time course of motion coherence is indicated above. Population output is unable to maintain high directionality as coherence drops. E) Population output at different time points during the trial. Time slices are shown at 400 ms intervals from start of first coherence segment. Left panel is for high coherence condition, middle panel for low coherence condition and right panel for decreasing coherence condition. Bottom panels show population response trajectories in TDR space. To the left are population trajectories in TDR space for high coherence condition. The responses are connected to form a high coherence contour. Middle panel shows trajectories for low coherence condition and the low coherence contour. Right panel shows decreasing coherence trajectories overlaid with high and low coherence contours. As coherence drops, population output is also not able to maintain high directionality. F) Mean activity level for an example stimulus direction for the three coherence conditions. Color coding as in D, black for high coherence, gray for low coherence and purple for decreasing coherence condition. Mean activity is similar across all three conditions. G) Population vector length for the same stimulus direction. High coherence stimulus shows high vector length indicating high directionality, low coherence condition lacks directionality and decreasing coherence conditions shows an initial high vector length which decreases with time.

**Supplementary figure 9:**
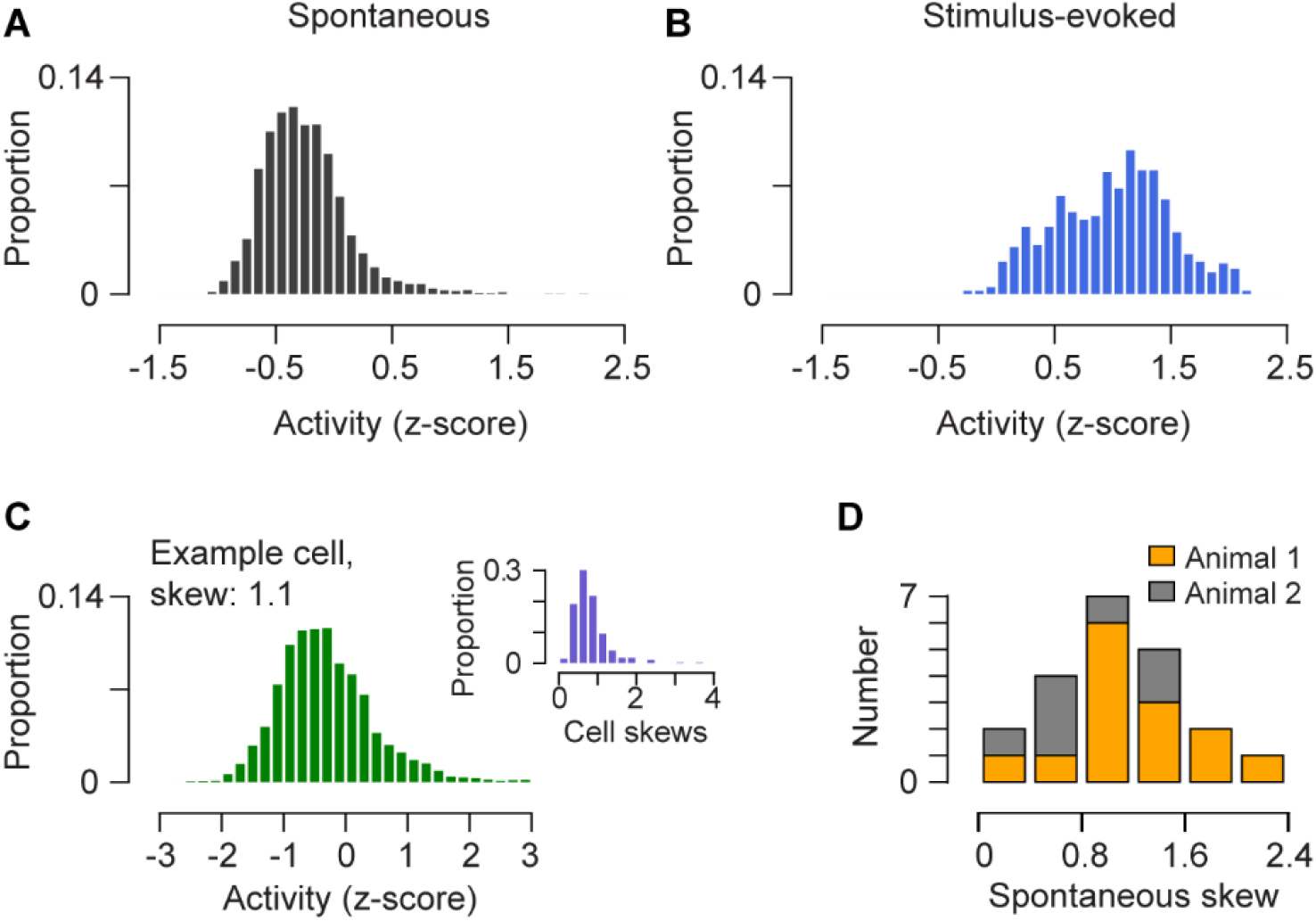
Activity distributions A) Spontaneous activity distribution of the mean population activity shown in fig 4. B) Stimulus-evoked activity distribution of the mean population activity during peak interval per trial for the same population. C) Spontaneous activity distribution of an example cell in the same population. The distribution has a skewness of 1.1. Inset shows the distribution of skews for spontaneous activity for all cells in the population. D) Distribution of spontaneous skews for mean population activity across different spontaneous sessions across the 2 animals.

**Supplementary figure 10:**
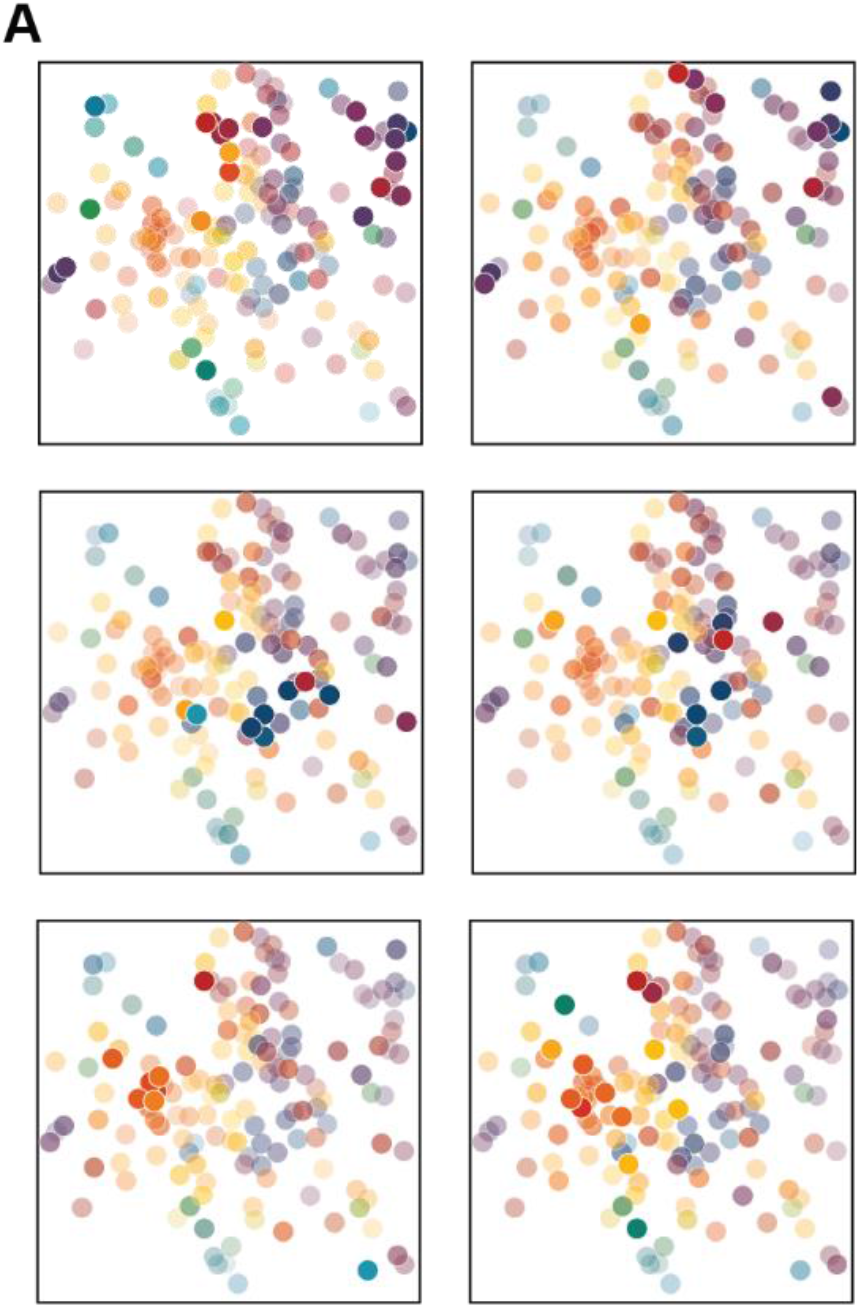
Spontaneously evoked activity patterns (A) Example patterns of activity observed in the absence of visual stimulation. The patterns occurred at 6 different time points. Intensity indicates the degree of activation, color indicates direction preference which is same as the map shown in figure 1D.

**Supplementary figure 11:**
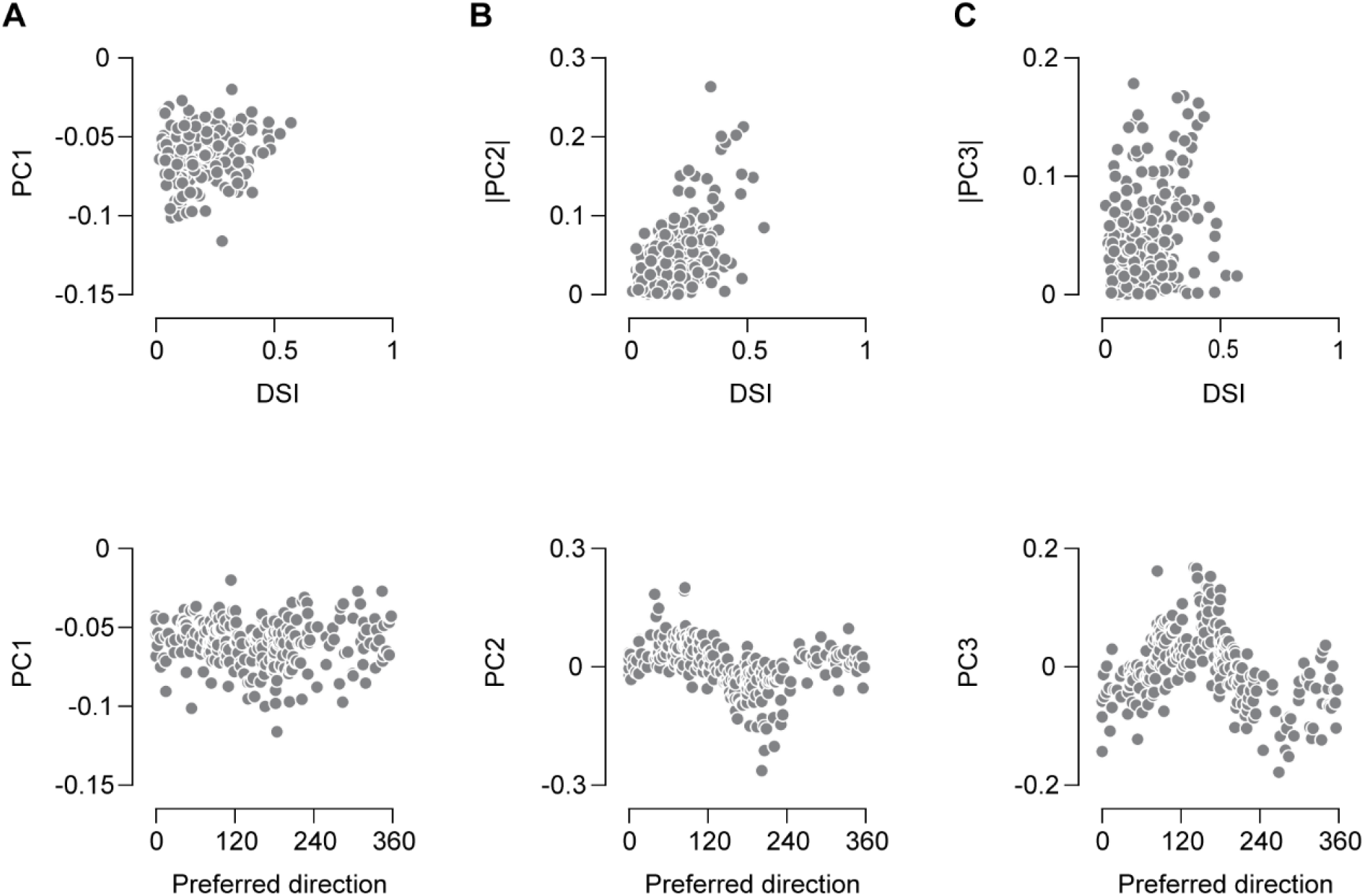
Characteristics of principal component weights. A-C) Top panels: Dependence of PC1 or the absolute value of PC 2 and 3 weights on DSI. Bottom panels: The relationship between of PCs 1-3 and preferred direction. There is a sinusoidal dependence between PC 2 and 3 weights and the neuronal preferred direction.

**Supplementary figure 12:**
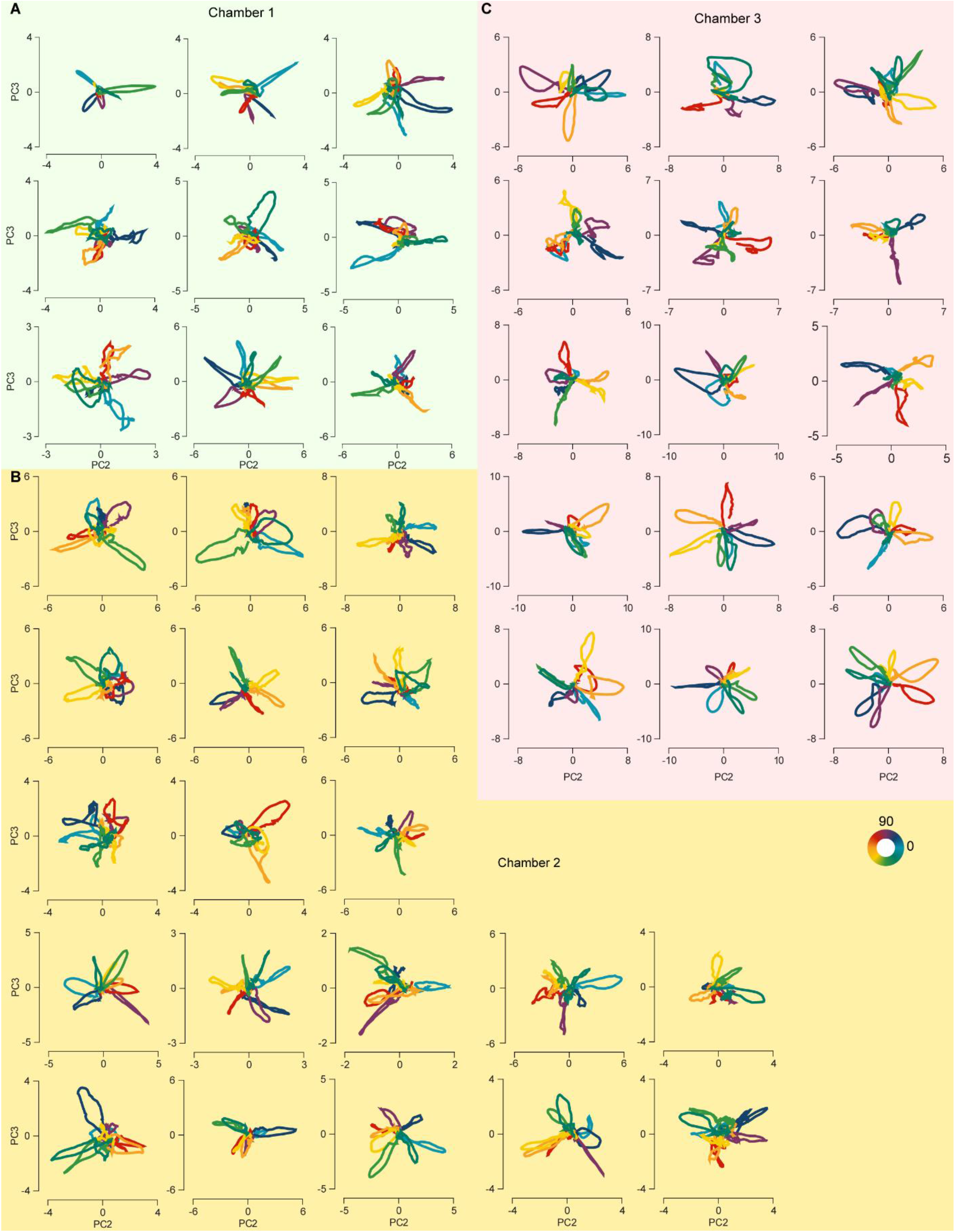
Examples of population trajectories in PC space showing direction selectivity across animals. A-C) Examples of stimulus-evoked population trajectories from three different chambers. Mean population response trajectories for the 8 stimulus directions are projected in their PC space. Each plot represents a different imaging session. Color coding of the trajectories represents the stimulus direction.

**Supplementary figure 13:**
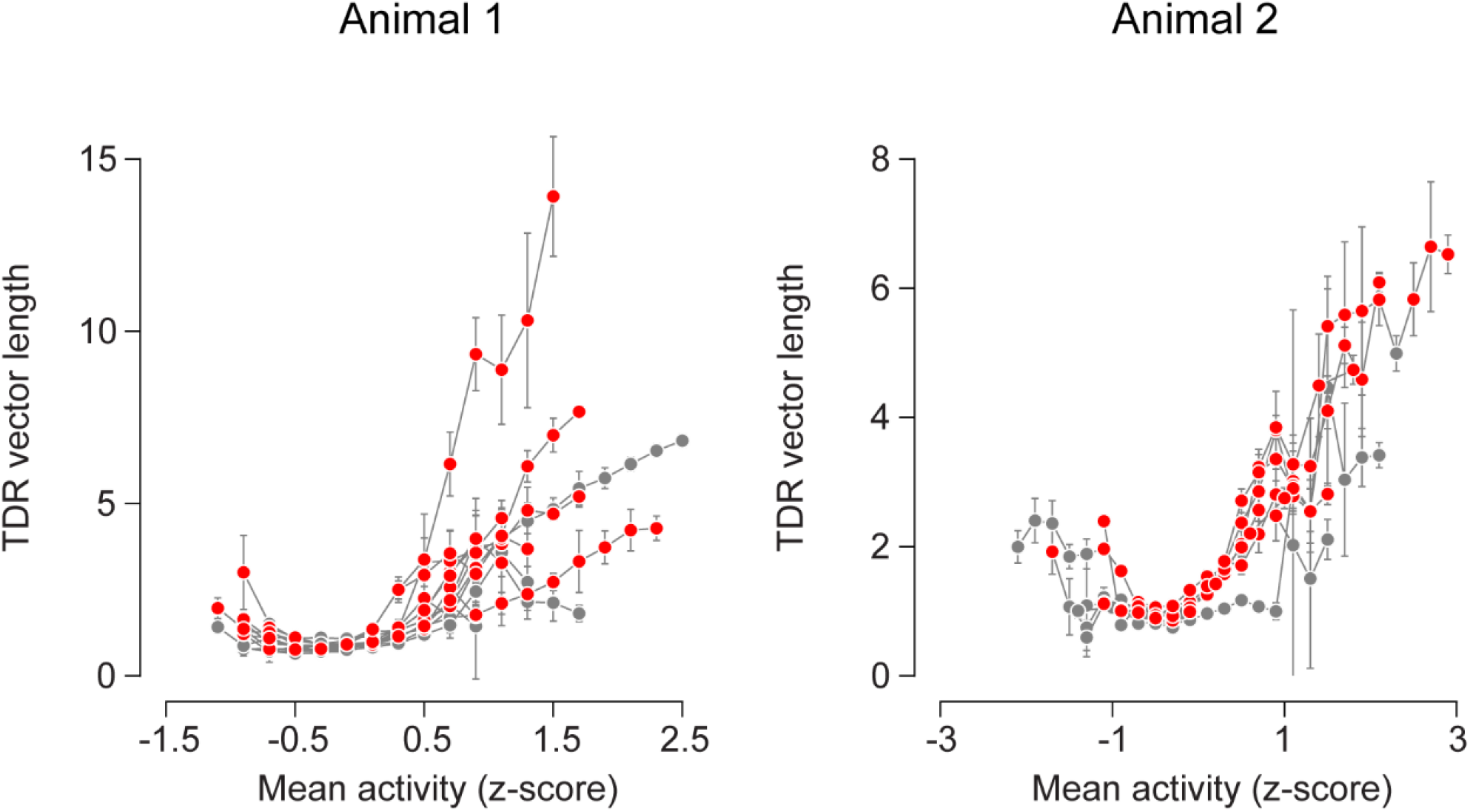
Activity dependent spontaneous population directionality. A,B) Dependence of output directionality on activity amplitude for different sessions. The vector length of the projection of spontaneous population activity in TDR space is plotted against the activity amplitude across sessions, in both animals. Each line represents one session. The observed median vector length values per activity bin are shown in red. The error bars indicate bootstrapped 95% confidence intervals. The corresponding values for shuffled data are shown in gray.

## Methods

All marmoset experiments were conducted with the approval of The University of Texas at Austin Institutional Animal Care and Use Committee.

One male and one female marmoset was used in the current study. One of the animals had MT chambers on both left and right side, while other had MT chamber only on the left side.

Surgical, virus injection and two-photon imaging procedures were similar to previous descriptions (*1, 2*).

### Surgery

Custom-made headpost (*3*) and chambers were affixed to the skull in a sterile anaesthetized procedure. Throughout the procedure, the body temperature was maintained at 36-37°C and the heart rate, SPO_2_ and CO_2_ were monitored. Animals were placed in stereotaxic frames, circular craniotomies were performed on the intended chamber location, over area MT, identified using stereotaxic coordinates, chambers and the headpost were placed and the dura was removed. An implant from dental acrylic was built around the headpost and chambers, covering the remaining exposed skull. The skin around the implant was affixed to the implant using Vetbond. The animals were then returned to the cages after recovery from anaesthesia.

### Chamber design

The chamber consisted of 4 parts. The outermost part of the chamber was a ring of height 1.6 mm and of diameters ranging from 5-7mm. This ring had 1 mm long thin feet that were inserted inside the skull following craniotomy. A second piece was a thin chamber nut (thickness 1.5 mm) that was screwed on the outside of the chamber ring and rested on top of the skull. This assembly was further sealed using Metabond (Parkell, New York). A removable imaging well was screwed on the inside of the chamber ring. The well consisted of a metal insert to which a coverglass was attached at the bottom. A thin cap (1 mm) was screwed on top of the chamber ring to close it.

### Virus injections

Virus injections were performed in a separated anaesthetized procedure. Animals were head-fixed, the head was disinfected and the procedure was performed under sterile conditions. The imaging well was removed from the chamber ring to physically access the cortex. The virus was injected using Nanoject II (Drummond Scientific) with pulled and beveled glass pipettes of tip diameters of 20-35 μm. rAAV constructs with GCaMP6f under the h56d promoter were used for these measurements (*1*). The glass pipette was filled with mineral oil and front loaded with the virus. Virus was injected at 23 nl/sec. Injections were made in multiple sites within the chamber at varying heights with each site receiving 500-1000 nl virus mix. The pipette was left in place for a 2-5 min before changing position. Injection spread was estimated using trypan blue dye diluted with the virus mix.

### Behavioral training and experimental control

After recovery from surgery, marmosets were habituated to head fixation and trained to fixate visual targets (*3*). Experimental control was provided by the Maestro software suite, which collected eye movement data, controlled visual stimulation, and provided juice reward (https://sites.google.com/a/srscicomp.com/maestro/).

### Acute recordings

The intracellular electrophysiology data presented in supplementary figure 4 was collected from recordings made in anaesthetized marmosets. The animals were anaesthetized on intravenous infusion of sufentanil and paralyzed using vecuronium. They were ventilated artificially. The vitals were continuously monitored and body temperature regulated at 37 degrees C. Once anaesthetized, area MT was located using stereotactic coordinates and a small craniotomy was performed to expose the brain surface.

### Stimulation

A screen subtending 48 by 38 degrees was placed 45 cm from the animal. Monitor frame rate was 100 Hz. To obtain direction selectivity of cells, full-field full coherence field of moving dots, speed 25 degrees/s, was presented to the animal at full contrast (dot density = 3.8 dots/deg^2^, dot size 0.4 degrees, black and white dots on gray background). The trial was composed of a blank period following which the moving stimulus was presented to the animal for a duration ranging from 700-1000 ms. Animal was free to move its eyes and received a small marshmallow juice reward at the end of each trial.

For spontaneous activity, the screen was set to gray background and there was no task and no reward. The spontaneous trials were interleaved between behavioral measurements.

For change direction experiment, a similar dot field was presented at full contrast, unless otherwise specified, and full coherence, moving along one direction for a certain period. The motion direction was suddenly switched to the second direction. Both directions were presented for equal amounts of time, which was either 500 ms or 750 ms each for imaging and 64 ms for extracellular electrophysiology. The direction change was either 90 or 135 degrees, varying for different sessions. Some sessions also presented an initial fixation target (2 deg by 2 deg) at the screen center. Fixation window was 3 degrees by 3 degrees and duration was 700 ms, with a grace period of 400 ms. Upon successful fixation, moving dot field was presented to the animal.

Animals were juice rewarded at the end of successful trial. Direction tuning was estimated from trials with only one motion direction presented per trial. Such trials were either interleaved with the change direction stimuli or were ran in a separate block.

For acute intracellular recordings, cells were presented with sinusoidal drifting gratings or plaids at their preferred directions and spatial frequencies.

### Two-photon imaging

Neuronal activity was measured using a custom-made two-photon microscope equipped with resonant mirrors for video rate sampling (30Hz) (*2*). Fluorescence was detected using standard PMTs (R6357, H7422PA-40 SEL, Hamamatsu, Japan) and amplified with a high-speed current amplifier (Femto DHPCA-100, Germany). Images were acquired using a 16x objective (Nikon N16XLWD-PF, Japan) from fields of view varying from 400 μm by 400 μm to 700 μm by 700 μm. Data were motion corrected using cross correlation (*4*). Any remaining frames where motion could not be corrected were manually marked for exclusion from analysis. Animals were imaged beginning 5-6 weeks post virus injection.

### Widefield imaging

Widefield maps were obtained by imaging using a 5x or 2.5x objective. For each stimulus trial, a prestimulus response was generated by averaging frames in the pre-stimulus interval and a stimulus response was generated by averaging frames during the peak response interval (10 to 24 frames after stimulus onset). The trial response was then computed by subtracting the prestimulus average from stimulus average. Trials at each stimulus direction were averaged to generate the mean response per direction. The obtained responses were smoothed using a 2-D Gaussian filter with a standard deviation of 1 to 3 pixels. For each pixel in the frame, the preferred direction was computed as the direction of the vector average of responses across presented directions.

### Intracellular recordings

Recordings were made with glass patch electrodes (5-10 MΩ) filled with potassium-gluconate based solution. Cells were recorded in whole-cell configuration. The spikes were removed by median filtering the raw membrane potential traces

### Imaging analysis

Cells were marked using a custom code in MATLAB and fluorescence values were extracted from the regions of interests. Slow drift, if any, was subtracted by computing a moving average over a 10 s period. ΔF/F was computed as follows:

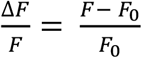

Where F is the raw fluorescence at each time point and F0 is the average baseline fluorescence. Responses were averaged across trials at each direction to give the mean response per direction. For the cell to be included in the analysis, the response at the maximum responsive direction had to be significantly different from baseline as measured using t-test (p: 0.05). In addition, the response at this maximally responsive direction also had to exceed a threshold, which varied between sessions from 0.1 to 0.25 ΔF/F. Traces were also median filtered with a 5^th^ order filter and smoothed with a moving average over 15 frames.

The direction selectivity index (DSI) was computed as follows:

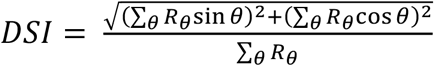

Standard error over the mean response was computed as follows:

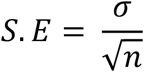

Where σ is the standard deviation of responses, and n is the number of trials.

The responses per trial were used to fit a von Mises direction tuning curve to each cell using least squares curve fitting. The form of the function is as below:

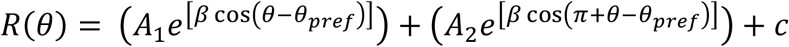

Where *R*(*θ*) is the response at direction θ, *A*_1_ is the maximum amplitude of the first peak, *A*_2_ is the maximum amplitude of the second peak 180 degrees away, β is the tuning width factor and *θ*_*pref*_ is the preferred direction of the cell and c is the amount of offset. The location of the peak of this fitted tuning curve is used as the preferred direction of each cell.

Distance between cells is calculated as the Euclidean distance between their measured x and y distance within the plane. For maps pooled across planes, the pairwise distance and difference in preferred directions are still computed only for within plane pairs.

The exponential fit to the data measuring dependence of preferred direction difference between cells with the distance between cells used the following equation:

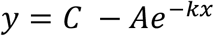

Where y is the fitted direction difference for x distance between cells, C is the saturation value, A is the start value and k is the decay space constant. The parameters were estimated using least squares curve fitting to individual data points.

The shuffled data for measuring this dependence was generated by keeping the same cell positions but shuffling their preferred directions.

Hypercolumn estimates were generated by fourier transform of the response maps obtained in wide field imaging.

### Principal Components Analysis (PCA)

Means and standard deviations were computed for each cell across all response traces across trials to be projected in the PC space. The data was then z-scored using these means and standard deviations. Covariance was computed over the mean responses for all directions. The eigenvectors of the PC space were then identified using singular value decomposition of the covariance matrix.

### Targeted dimensionality reduction (TDR)

To directly estimate the relevant population response space explaining motion related variance, we used TDR (adapted from (*5*)). The original data was converted to reduced dimensionality by keeping principal dimensions 2-10. The response at each time point in the trial could be described using the following equation:

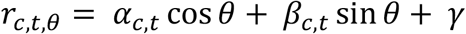

Where *r* is the response of cell c at time t in the trial for stimulus θ. The regression coefficients α and β define the direction subspace and γ is offset. The regression coefficients were estimated by taking the Fourier transform of the response vector across all directions at each time in the trial. The time-independent regression coefficients were identified as the coefficients at time point t in the trial such that:

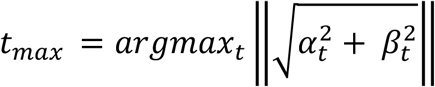

These time independent regression coefficients were then orthogonalized using QR decomposition. The resulting vectors were used to project the data in the direction subspace.

### TDR vector length was computed as follows

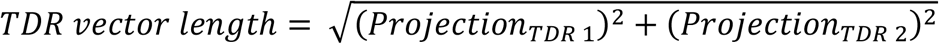

Error bars in Fig 4J on binned vector lengths as a function of activity were 95% bootstrapped confidence intervals on the median vector length.

### Extracellular electrophysiology analysis

Single unit records from macaque area MT were measured as in (6). The response of each single unit at its preferred direction was used to estimate the response decay timing. Cell responses were normalized by dividing each cell’s response by its sustained firing rate during stimulation at preferred direction. Response latency was identified as the time it took for the mean baseline subtracted response to exceed 5% of its peak value, following motion onset. Response at this latency following motion offset was used as starting point for decay analysis. An exponential decay was fit to the cell’s response. The form of the function is as follows:

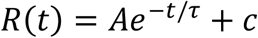

Where R(t) is the response at interval t following motion offset and response latency. A is the amplitude of response at initial point, τ is the decay time constant in ms and c is the offset.

Pre and post-stimulus activity was averaged from a 64 ms interval. Prestimulus interval was 64 ms before the start of the stimulus. Post-stimulus interval started at 188 ms following end of stimulus.

### Intracellular electrophysiology analysis

Responses were median filtered with filter 30 ms to remove spikes. The mean response was baseline subtracted. Sustained response was calculated by averaging response in the last 1s before stimulus offset. The mean response was then normalized by dividing by the sustained response. The response to stimulus offset was fitted with the same exponential decay as above.

### Change direction analysis

For both imaging and electrophysiology measures, neural responses were z-scored using the mean and standard deviation across repeats across different conditions. For the change direction stimulus, the responses were binned into direction bins for each condition, averaging the responses of cells with preferred direction within one bin. The bins were then rotated per condition so that 1^st^ and 2^nd^ stimulus direction were aligned for all conditions. Then the responses were averaged across conditions.

For the partial correlation analysis, mean population response to change direction conditions was binned in direction bins (averaging cells with direction preference within each bin to generate the bin response) and time bins. The response in each time bin was fitted with two fit functions, either to generate the switch or the sweep prediction. The switch prediction was generated as fit to the response was with sum of two gaussians separated by the degrees of difference between the two stimulus directions (90 or 135). The form of the function is as below:

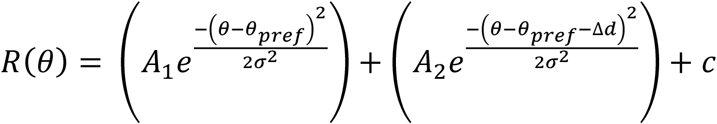

Where R is the response in direction bin θ, A_1_ and A_2_ are the amplitudes for the two peaks, σ is the width of the Gaussian, Δd is the difference in the two stimulus directions and c is the offset. The width of the Gaussian was set to be a constant ±1, making σ to effectively be a set parameter instead of a free parameter. This is done so that the number of free parameters (four) is same for the switch and sweep fits.

The sweep prediction was generated as the fit to response with a single Gaussian. The form of the function is below, with parameters same as above.

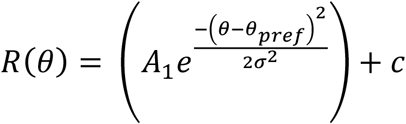

The correlation values were computed as c_switch_ for correlation between the data and switch prediction, c_sweep_ for correlation between the data and the sweep prediction, c_swith-sweep_ for correlation between the switch and sweep predictions. The partial correlation values were computed as follows:

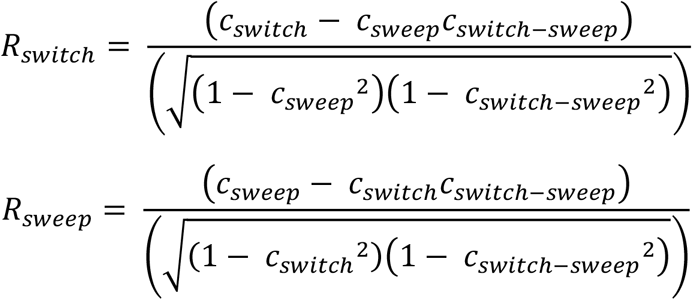

The population output for different time slices was fit with sum of two gaussians separated by 135 degrees, the amplitude of second gaussian could be 0, and another free parameter indicated the response offset across all directions. Standard deviation of the gaussians was set to be constant.

For electrophysiology analysis in supplementary figure 6, 33 single units were used. From each unit, 8 cell profiles were generated by shifting the cell tuning in 45 degrees step. This generated a population of 264 cells. The predicted rate shown in supplementary figure 6C was conservatively estimated as max of response to stimulus 1 and response to stimulus 2 shifted in time by stimulus 1 duration. Rate is calculated as mean firing within 192 ms following start of stimulus 1. Even with the conservative estimate, the observed response was lower than predicted, defying expectation of the ring model.

### Change coherence analysis

Responses of cells were z-scored using the mean and standard deviation across repeats across different conditions. For the change direction stimulus, the responses were binned into direction bins for each condition. The bins were then rotated per condition so that stimulus direction was aligned for all conditions. Then the responses were averaged across conditions.

### Numerical simulations

For demonstrating the four model regime predictions under different scenarios, simulations were ran for following connectivity parameters. J_0_^E^ was always set to 0 and J_0_^I^ to 0.7:

| | $J_1^E$ | $J_1^I$ |
| --- | --- | --- |
| f-NA | 0.15 | 0.0937 |
| s-NA | 1.8 | 1.1250 |
| CA | 4.5 | 2.8125 |
| BA | 4.2 | 4.2 |

BA corresponds to the transient amplification regime in the analytical derivations, f-NA and s-NA are in the homogenous regime and CA is in the marginal regime.

The model network consisted of 512 excitatory cells and 512 inhibitory cells with preferred directions spaced evenly across E cells and I cells. Connectivity weight from an excitatory cell j to cell i is as follows:

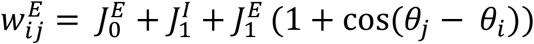

Where θ is the direction preference of the cell.

Similarly, connectivity weight from inhibitory cell j to cell i is:

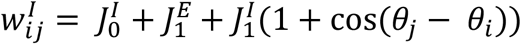

To simulate single population response, weights were defined as

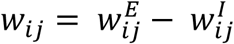

Simulations were run at 0.1 ms time step. Input at each time step was generated from a sum of local input, external input and noise input.

Local input was modeled as *Ws* where W is the weight matrix of the population and s is the response matrix. External input was modeled differently based on the required predictions. The general form of the external input was

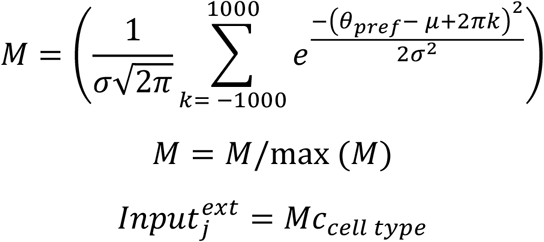

Where c_celltype_ was set based on whether cell was E or I. σ controlled the width of the stimulus and was set based on context. Θ_pref_ is the cell’s preferred direction, μ is the stimulus direction.

If noise input was used, a multiplicative noise was independently added to each cell. The noise was drawn from a normal distribution of mean 0 and standard deviation of 0.5. Additionally, a constant noise input, when used, was set to a value of 1. The sum of all inputs was rectified (*7, 8*) and passed through a saturating non-linearity (output was capped at 10). Each iteration of network output was simulated using the rate equation and decay constant was set to 20 ms.

The numerical response maps in supplementary figure 3 were generated by stimulating across range of connectivity parameters. Each trial was 3 s long, external input was on for first half of the trial with μ set to zero and σ set to 1.2. No noise was used in this context. J_0_^I^ of 0.2 was used here. The numerical amplification was defined as selectivity of the output divided by selectivity of the input. Selectivity was defined as

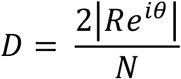

Where R is response of each model cell, θ is its direction preference, and N is the total number of cells in the network.

To simulate the change direction predictions, network was provided input at first stimulus direction in the first half of the trial. In the second half of the trial, the stimulus direction was shifted to the second direction. The input was defined as above with c set to 2 for both cell types, stimulus direction μ as 90 for stimulus 1 and 225 for stimulus 2, σ was set to 1.2. No noise was used in this context. Each trial was 1s long.

To simulate the change coherence predictions, network was provided input at a single direction of 180. Each trial was divided into 4 segments of equal duration, total length of the trial being 2s long. For the external stimulus, c was set to 2k for both cell types, k being a scaling factor changing with coherence levels, μ was 180, σ was set to 1.2. k was set to 1, 0.7, 0.225 and 0.025, to match the stimulus coherence levels of 100, 70, 22.5 and 2.5%. To keep the area under the stimulus curve (to replicate equal motion for all coherence levels) same for all coherence levels, additional offsets were added to coherence levels less than 100. No noise was used in this context.

To simulate spontaneous activity, no external input was used. Every cell received a noise drawn from a normal distribution of mean 0 and standard deviation of 0.5. Additionally, a constant noise input, of value 1 was added to all cells. The output angle of this activity was computed by estimating the x and y components after summing each cell’s activity multiplied by its direction preference and dividing this value by the sum of responses. The vector length of the output at each time step was calculated as follows:

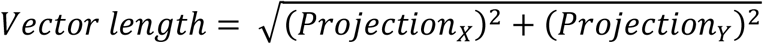

The corresponding activity level was computed as the mean activity across all model cells. This activity was binned and the vector lengths for time points within each bin were averaged to give the mean vector length value at each activity bin. The error bars were computed as the standard deviation over these vector lengths.

Analytical derivations are described in detail later. Briefly, the plots in figure 2 D-F are based on analytical derivations. Amplification (see description of gain in analytical derivations is defined as: 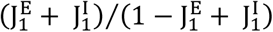.

Scaled time constant is defined as: 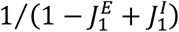.

## Cortical selective amplification models

### 1 Network dynamics

Our general equation for a recurrent cortical network of *N* excitatory (E) and *N* inhibitory (I) neurons (in vector-matrix notation) is:

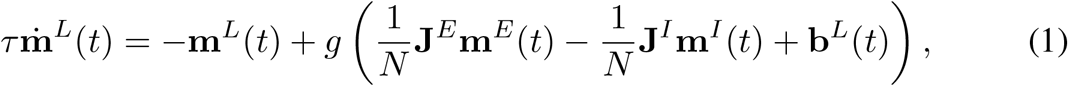

where *g* is the nonlinear neural activation function, **m**^*L*^ is the vector of activations of neurons in population *L*, with *L* = {*E, I*} indexing the E, I populations, respectively. 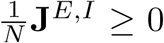 are the weights from the excitatory and inhibitory neurons to all neurons.

Writing out the indices explicitly, the equation is:

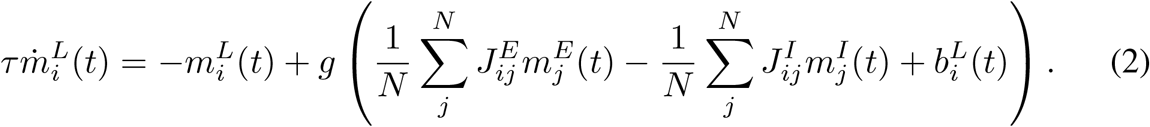

In the continuum limit (*N* →, *i* → *θ*, and 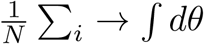), we obtain the following neural field model:

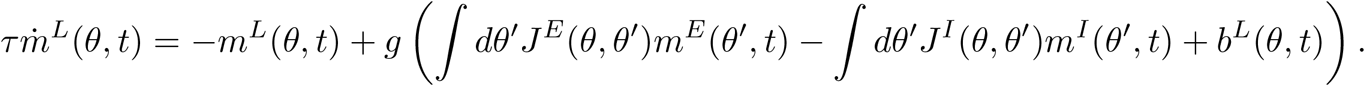

If we assume that the weights depend only on the difference in the indices of the neurons, *J*(*θ, θ*^*′*^) = *J*(|*θ* − *θ*^*′*^|) (translation or rotation invariance), as we will assume everywhere, then

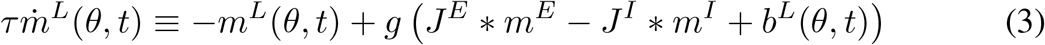

where ∗ refers to the convolution operation.

### 2 Network architecture

For all models presented in the main manuscript, we consider a unified framework with a common architecture. The different models can then be obtained by considering different parametric regimes of the network. The network has structured lateral connectivity, in which neurons are assigned a preferred angle, *θ* ∈ [0, 2*π*], and both E and I neurons maximally project to similarly tuned neurons 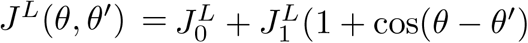, where 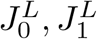 are the amplitudes of the spatially untuned and tuned components, respectively, of the *L*th population. The external or feedforward input 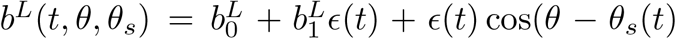 similarly consists of a spatially untuned component 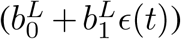 and tuned component (*ϵ* (*t*) cos(*θ* − *θ*_*s*_(*t*))), with 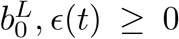 and where *θ*_*s*_ is the stimulus direction. The tuned input may be time-dependent, modeled by a time-dependent *ϵ* or *θ*_*s*_(*t*), depending on whether the amplitude or angle are time-varying. This connectivity and input structure are similar to the ring model of [1], generalized to two neural populations (E,I).

### 3 Linearized dynamics

We seek to determine the stability of a uniform fixed point 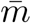 when the tuned component of the input is 0. We do so by considering how the dynamics of Equation 3 governs small deviations 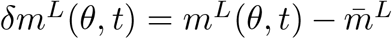 away from this state. Rewriting Equation 3 after assuming a constant input 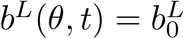, we have:

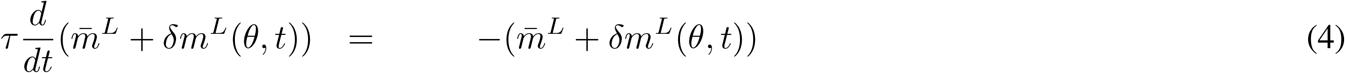

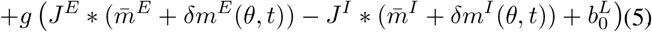

Using the assumption that 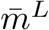 is a fixed point, and Taylor expanding the nonlinear function *g*() to first order around 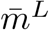, we have that

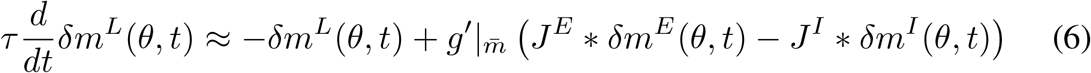

where we have ignored terms of higher order in *δm*^*L*^.

Under our assumption that the weights are translation-invariant (*J* (*θ, θ*^*′*^) = *J* (|*θ θ*^*′*^|)), the eigenvectors of the weights are Fourier modes, and thus the recurrently coupled network’s dynamics naturally decompse into the independent dynamics of a set of Fourier modes. Fourier transforming Eq. 6 (applying *∫ dθe*^*ikθ*^ to all sides), we get:

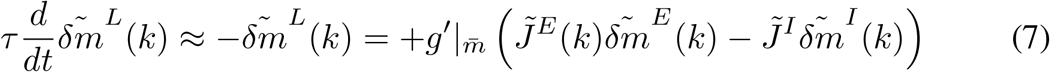

where 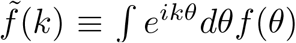 refers to the Fourier transform of *f*. Note that under the Fourier transformation, the recurrently coupled dynamics have been replaced by a set of separate equations for each wavenumber *k*, shown in Equation 7. In our model, 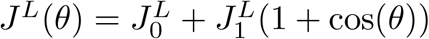 so

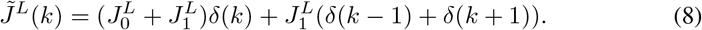

### 4 Single-population system and regimes

If all neurons were of a single type with combined weights *J*^*E*^ − *J*^*I*^, rather than of types E, I separately, the linearized dynamics of Eq. 7 would be given by:

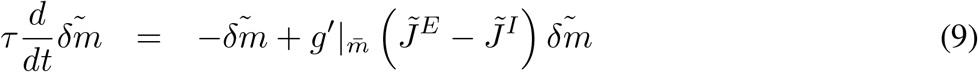

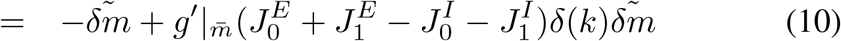

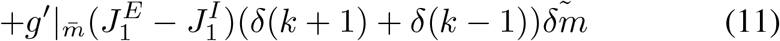

The stability of the uniform mode of activation (the DC or *δ*(*k*) mode) in this linear dynamical system is simply determined by 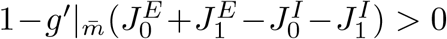. Since we are considering threshold-linear units, 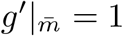, hence the condition for stability of the uniform mode in the single-population case with rectified linear activation functions is simply 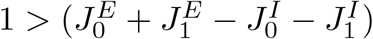 or

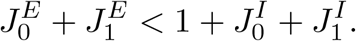

This condition is required for stability of the system, in both the marginal (continuous attractor or CA) and the homogeneous (single stable fixed point) regimes. Next, for the CA regime, the network must exhibit an emergent, self-sustained patterned state with an activity bump. This requires instability of the *k* = 1 mode; thus, the condition for the CAregime is that 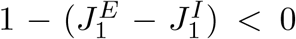, or in other words, the existence of CAdynamics in the single population case requires:

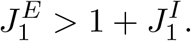

The single population homogeneous regime, by contrast, has:

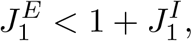

with the degree of selective amplification (gain factor) given by 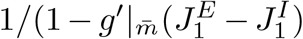 (Eq. 11), which in the case of threshold-linear neurons becomes 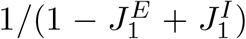. Maximal selective amplification within the homogeneous regime is therefore given by 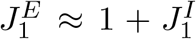, which is a finely-tuned regime right at the boundary of the homogeneous and CAregimes; simultaneously, to minimize DC amplification, 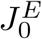 should be minimized, so 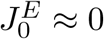.

### 5 System with separate E,I-populations

Inserting expressions for the Fourier-transformed weights and collecting like terms in Eq. 7 for the E and I populations, we have:

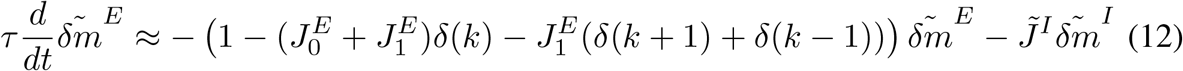

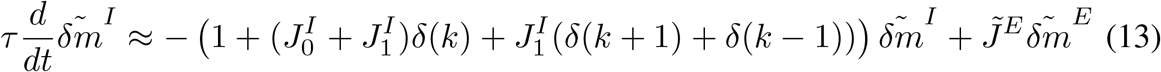

where we have used 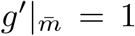 above. Note that the inhibitory population is always stable since the prefactor of 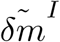 on the right hand side of the second equation is always negative (assuming 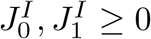). For the *k* = 0 mode, the equations are:

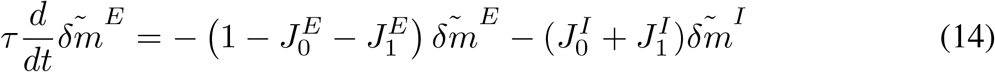

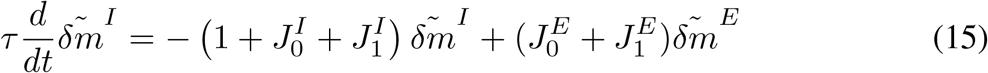

which can be summarized as 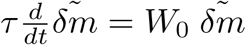 where

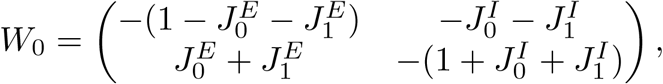

while for *k* = *±*1, they are:

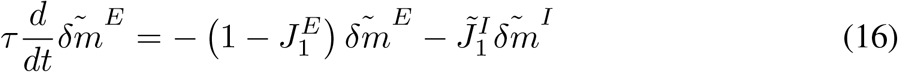

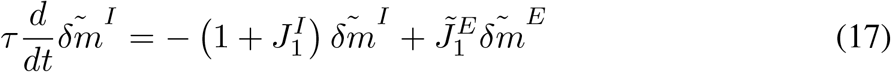

which can be summarized as 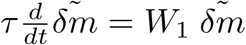 where

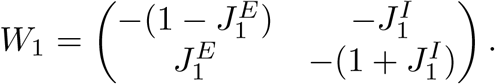

The dynamics are stable if both eigenvalues of each of *W*_0_ and *W*_1_ are negative. One eigenvalue of each matrix is −1 and thus automatically negative, while the second of each is given by 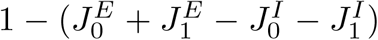 and 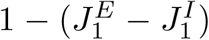, respectively. Thus, the conditions for stability in the two-population case are:

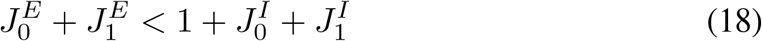

and

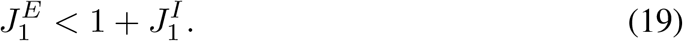

Note that these conditions on stability are identical to those in the single-population case. However, the two-population network exhibits a different dynamics than the single-population case, arising from the fact that the matrices *W*_0_, *W*_1_, which control the uniform and tuned responses of the network, are non-normal. The dynamics of a normal network can always be decomposed into independent eigenmodes by diagonalizing the weight matrix. The dynamics of a system with a non-normal interaction matrix typically cannot be decomposed in this way because non-normal matrices are typically only reducible by orthogonal transformation to upper-triangular rather than diagonal form.

Any real invertible matrix can be written in triangular form by an orthogonal transformation; for instance for *W*_1_ there is some orthogonal matrix *O* such that *O*^*T*^*W*_1_*O* = *L*_1_, where *L*_1_ is an upper-traingular matrix. The interpretation of this mathematical fact in network terms is that a non-normally interacting recurrent network is equivalent to some feedforward network with interaction matrix *L*. The matrix *O* can be found by selecting one of its columns to be one of the normalized eigenvectors of *W*_1_, which in our case includes the vector 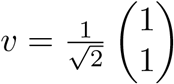. The other column is then a normalized vector orthogonal to the selected eigenvector. In this way, we find that 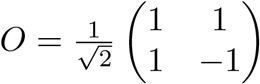, and therefore that

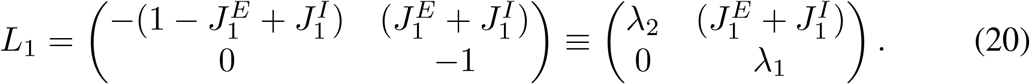

where *λ*_1,2_ are the two eigenvalues of *W*_1_. In other words, the transformed network dynamics for the tuned mode is given by the feedforward matrix *L*_1_ acting on 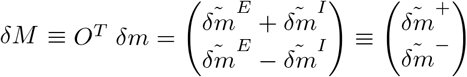. Thus, we have:

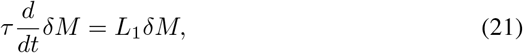

which, written out in full form, yields:

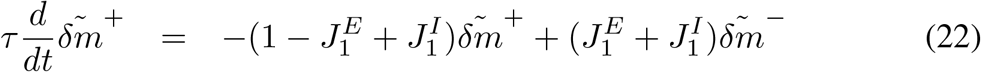

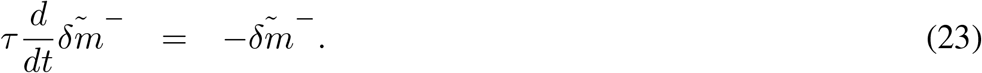

In other words, any inputs fed into the difference mode 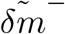 get scaled by a feed-forward gain factor 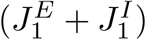 and funnel into the sum mode. The sum and difference and modes are themselves stable, decaying with time-constants 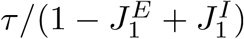 and *τ*, respectively. Note that when 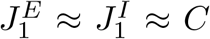 (near-balanced), both modes are fast, with a decay that is as rapid as the single-neuron time-constant.

The total gain in the sum mode for inputs fed into the differnce mode is given by 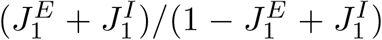. Compare this to the gain from the single-population (normal amplification) regime, which is 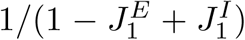; thus we see that the total gain of the sum mode consists of a normal amplification component (the denominator), and a non-normal or feedforward amplification of differential inputs to the E and I populations (the numerator). When this quantity is positive and large, the sum mode will display a large (but transient) amplification. Thus, a network that is nearly balanced with strong tuned excitation and strong tuned inhibition is in the tuned transient amplification regime [2]. The coexistence of fast response and large amplification is in contrast to the homogoneous and CAregimes of the single-population network, in which amplification goes hand-in-hand with slow dynamics: The amplfication factor in the homogeneous and CAregimes is 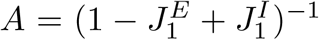 and the corresponding time-constant of the dynamics is *Aτ*. This means that in the limit of maximal amplification within the normal amplification regime, the time-constant of the tuned mode diverges.

## References

1. Sclar, G. & Freeman, R. D. Orientation Selectivity in the Cat’s Striate Cortex is Invariant with Stimulus Contrast*. Exp Brain Res vol. 46 (1982).

2. Albrecht, D. G. & Hamilton, D. B. Striate cortex of monkey and cat: Contrast response function. J. Neurophysiol. 48, 217–237 (1982).

3. Skottun, B. C., Bradley, A., Sclar, G., Ohzawa, I. & Freeman, R. D. The effects of contrast on visual orientation and spatial frequency discrimination: A comparison of single cells and behavior. J. Neurophysiol. 57, 773–786 (1987).

4. Britten, K. H., Newsome, W. T., Shadlen, M. N., Celebrini, S. & Movshon, J. A. A relationship between behavioral choice and the visual responses of neurons in macaque MT. Visual Neuroscience vol. 13 (1996).

5. Anderson, J. S., Carandini, M. & Ferster, D. Orientation tuning of input conductance, excitation, and inhibition in cat primary visual cortex. J. Neurophysiol. 84, 909–926 (2000).

6. Hubel, D. H. & Wiesel, T. N. Receptive fields, binocular interaction and functional architecture in the cat’s visual cortex. J. Physiol. 160, 106–154 (1962).

7. Ferster, D. & Lindström, S. An intracellular analysis of geniculo-cortical connectivity in area 17 of the cat. J. Physiol. 342, 181–215 (1983).

8. Douglas, R. J., Koch, C., Mahowald, M., Martin, K. A. C. & Suarez, H. H. Recurrent excitation in neocortical circuits. Science 269, 981–985 (1995).

9. Chung, S. & Ferster, D. Strength and orientation tuning of the thalamic input to simple cells revealed by electrically evoked cortical suppression. Neuron 20, 1177–1189 (1998).

10. Chance, F. S., Nelson, S. B. & Abbott, L. F. Complex cells as cortically amplified simple cells. Nat. Neurosci. 2, 277–282 (1999).

11. Li, L. Y., Li, Y. T., Zhou, M., Tao, H. W. & Zhang, L. I. Intracortical multiplication of thalamocortical signals in mouse auditory cortex. Nat. Neurosci. 16, 1179–1181 (2013).

12. Lien, A. D. & Scanziani, M. Tuned thalamic excitation is amplified by visual cortical circuits. Nat. Neurosci. 16, 1315–1323 (2013).

13. Li, Y. T., Ibrahim, L. A., Liu, B. H., Zhang, L. I. & Tao, H. W. Linear transformation of thalamocortical input by intracortical excitation. Nat. Neurosci. 16, 1324–1330 (2013).

14. Zerlaut, Y., Zucca, S., Panzeri, S. & Fellin, T. The Spectrum of Asynchronous Dynamics in Spiking Networks as a Model for the Diversity of Non-rhythmic Waking States in the Neocortex. Cell Rep. 27, 1119-1132.e7 (2019).

15. Peron, S. et al. Recurrent interactions in local cortical circuits. Nature 579, 256–259 (2020).

16. Dubner, R. & Zeki, S. M. Response properties and receptive fields of cells in an anatomically defined region of the superior temporal sulcus in the monkey. Brain Res. 35, 528–532 (1971).

17. Rosa, M. G. P. & Elston, G. N. Visuotopic organisation and neuronal response selectivity for direction of motion in visual areas of the caudal temporal lobe of the marmoset monkey (Callithrix jacchus): Middle temporal area, middle temporal crescent, and surrounding cortex. J. Comp. Neurol. 393, 505–527 (1998).

18. Solomon, S. S. et al. Visual motion integration by neurons in the middle temporal area of a New World monkey, the marmoset. J. Physiol. 589, 5741–5758 (2011).

19. Lui, L. L. & Rosa, M. G. P. Structure and function of the middle temporal visual area (MT) in the marmoset: Comparisons with the macaque monkey. Neuroscience Research vol. 93 (2015).

20. Mehta, P. et al. Functional Access to Neuron Subclasses in Rodent and Primate Forebrain. Cell Rep. 26, 2818-2832.e8 (2019).

21. Ohki, K., Chung, S., Ch’ng, Y. H., Kara, P. & Reid, R. C. Functional imaging with cellular resolution reveals precise microarchitecture in visual cortex. Nature 433, 597–603 (2005).

22. Albright, T. D., Desimone, R. & Gross, C. G. Columnar organization of directionally selective cells in visual area MT of the Macaque. J. Neurophysiol. 51, (1984).

23. Hubel, D. H. & Wiesel, T. N. Sequence regularity and geometry of orientation columns in the monkey striate cortex. J. Comp. Neurol. 158, 267–293 (1974).

24. Goldberg, J. A., Rokni, U. & Sompolinsky, H. Patterns of ongoing activity and the functional architecture of the primary visual cortex. Neuron 42, 489–500 (2004).

25. Murphy, B. K. & Miller, K. D. Balanced Amplification: A New Mechanism of Selective Amplification of Neural Activity Patterns. Neuron 61, 635–648 (2009).

26. Van Vreeswijk, C. & Sompolinsky, H. Chaos in neuronal networks with balanced excitatory and inhibitory activity. Science 274, 1724–1726 (1996).

27. Ben-Yishai, R., Hansel, D. & Sompolinsky, H. Traveling waves and the processing of weakly tuned inputs in a cortical network module. J. Comput. Neurosci. 4, 57–77 (1997).

28. Buzsáki, G. & Mizuseki, K. The log-dynamic brain: How skewed distributions affect network operations. Nat. Rev. Neurosci. 15, 264–278 (2014).

29. Tan, A. Y. Y., Chen, Y., Scholl, B., Seidemann, E. & Priebe, N. J. Sensory stimulation shifts visual cortex from synchronous to asynchronous states. Nature 509, 226–229 (2014).

30. Arieli, A., Sterkin, A., Grinvald, A. & Aertsen, A. Dynamics of ongoing activity: Explanation of the large variability in evoked cortical responses. Science 273, 1868–1871 (1996).

31. Kenet, T., Bibitchkov, D., Tsodyks, M., Grinvald, A. & Arieli, A. Spontaneously emerging cortical representations of visual attributes. Nature 425, 954–956 (2003).

32. Luczak, A., Barthó, P. & Harris, K. D. Spontaneous Events Outline the Realm of Possible Sensory Responses in Neocortical Populations. Neuron 62, 413–425 (2009).

33. Berkes, P., Orbán, G., Lengyel, M. & Fiser, J. Spontaneous cortical activity reveals hallmarks of an optimal internal model of the environment. Science 331, 83–87 (2011).

34. Omer, D. B., Fekete, T., Ulchin, Y., Hildesheim, R. & Grinvald, A. Dynamic patterns of spontaneous ongoing activity in the visual cortex of anesthetized and awake monkeys are different. Cereb. Cortex 29, 1291–1304 (2019).

35. Davis, Z. W., Muller, L., Martinez-Trujillo, J., Sejnowski, T. & Reynolds, J. H. Spontaneous travelling cortical waves gate perception in behaving primates. Nature 587, 432–436 (2020).

36. Mante, V., Sussillo, D., Shenoy, K. V. & Newsome, W. T. Context-dependent computation by recurrent dynamics in prefrontal cortex. Nature 503, 78–84 (2013).

37. Stringer, C. et al. Spontaneous behaviors drive multidimensional, brainwide activity. Science 364, (2019).

38. Ozeki, H., Finn, I. M., Schaffer, E. S., Miller, K. D. & Ferster, D. Inhibitory Stabilization of the Cortical Network Underlies Visual Surround Suppression. Neuron 62, 578–592 (2009).

39. Ahmadian, Y., Rubin, D. B. & Miller, K. D. Analysis of the stabilized supralinear network. Neural Comput. 25, 1994–2037 (2013).

40. Hennequin, G., Vogels, T. P. & Gerstner, W. Optimal control of transient dynamics in balanced networks supports generation of complex movements. Neuron 82, 1394–1406 (2014).

41. Rubin, D. B., VanHooser, S. D. & Miller, K. D. The stabilized supralinear network: A unifying circuit motif underlying multi-input integration in sensory cortex. Neuron 85, 402–417 (2015).

42. Hennequin, G., Ahmadian, Y., Rubin, D. B., Lengyel, M. & Miller, K. D. The Dynamical Regime of Sensory Cortex: Stable Dynamics around a Single Stimulus-Tuned Attractor Account for Patterns of Noise Variability. Neuron 98, 846-860.e5 (2018).

43. Cohen, M. R. & Maunsell, J. H. R. Attention improves performance primarily by reducing interneuronal correlations. Nat. Neurosci. 12, 1594–1600 (2009).

44. Mitchell, J. F., Sundberg, K. A. & Reynolds, J. H. Spatial Attention Decorrelates Intrinsic Activity Fluctuations in Macaque Area V4. Neuron 63, 879–888 (2009).

45. McGinley, M. J. et al. Waking State: Rapid Variations Modulate Neural and Behavioral Responses. Neuron 87, 1143–1161 (2015).

46. McGinley, M. J., David, S. V. & McCormick, D. A. Cortical Membrane Potential Signature of Optimal States for Sensory Signal Detection. Neuron 87, 179–192 (2015).

47. Zhang, K. Representation of spatial orientation by the intrinsic dynamics of the head-direction cell ensemble: A theory. J. Neurosci. 16, 2112–2126 (1996).

## References

1. P. Mehta, L. Kreeger, D. C. Wylie, J. J. Pattadkal, T. Lusignan, M. J. Davis, G. F. Turi, W. K. Li, M. P. Whitmire, Y. Chen, B. L. Kajs, E. Seidemann, N. J. Priebe, A. Losonczy, B. V. Zemelman, Functional Access to Neuron Subclasses in Rodent and Primate Forebrain. Cell Rep. 26, 2818-2832.e8 (2019).

2. J. J. Pattadkal, G. Mato, C. van Vreeswijk, N. J. Priebe, D. Hansel, Emergent Orientation Selectivity from Random Networks in Mouse Visual Cortex. Cell Rep. 24, 2042-2050.e6 (2018).

3. J. F. Mitchell, N. J. Priebe, C. T. Miller, Motion dependence of smooth pursuit eye movements in the marmoset. J. Neurophysiol. 113, 3954–3960 (2015).

4. M. Guizar-Sicairos, S. T. Thurman, J. R. Fienup, Efficient subpixel image registration algorithms. Opt. Lett. 33 (2008), doi:10.1364/ol.33.000156.

5. V. Mante, D. Sussillo, K. V. Shenoy, W. T. Newsome, Context-dependent computation by recurrent dynamics in prefrontal cortex. Nature. 503, 78–84 (2013).

6. N. J. Priebe, S. G. Lisberger, Constraints on the Source of Short-Term Motion Adaptation in Macaque Area MT. II. Tuning of Neural Circuit Mechanisms. J Neurophysiol. 88, 370–382 (2002).

7. R. Ben-Yishai, R. Lev Bar-Or, H. Sompolinsky, Theory of orientation tuning in visual cortex. Proc. Natl. Acad. Sci. U. S. A. 92, 3844–3848 (1995).

8. R. Ben-Yishai, D. Hansel, H. Sompolinsky, Traveling waves and the processing of weakly tuned inputs in a cortical network module. J. Comput. Neurosci. 4, 57–77 (1997).

## References

[1] R Ben-Yishai, RL Bar-Or, and H Sompolinsky. Theory of orientation tuning in visual cortex. Proc Natl Acad Sci U S A, 92(9):3844–8, Apr 25 1995.

[2] Brendan K Murphy and Kenneth D Miller. Balanced amplification: a new mechanism of selective amplification of neural activity patterns. Neuron, 61(4):635–48, Feb 2009.

